# A Dual-Function Guanidinium Scaffold that Couples Copper Sequestration with Redox Protection in Wilson disease

**DOI:** 10.64898/2026.06.11.731572

**Authors:** Raviranjan Pandey, Arpan N Roy, Sandip Sarkar, Krishanu Dutta, Debosmita Bhattacharya, Akhilesh Jaiswar, Tamal Ghosh, Kalyan Goswami, Chinmoy Patra, Amitava Das, Arnab Gupta

**Affiliations:** Department of Biological Sciences, Indian Institute of Science Education and Research Kolkata, Mohanpur, West Bengal 741246, India; Department of Chemical Sciences, Indian Institute of Science Education and Research Kolkata, Mohanpur, West Bengal 741246, India; Agharkar Research Institute, G. G. Agarkar Road, Pune, Maharashtra 411004, India; Department of Biochemistry, All India Institute of Medical Sciences (AIIMS), Kalyani, West Bengal 741245, India

**Keywords:** Copper homeostasis, Wilson disease, Copper reactivity, Guanidium scaffold, Redox modulation, Metal dyshomeostasis

## Abstract

Wilson disease (WD) is caused due to mutations in the copper ATPase gene ATP7B, that result in accumulation of labile copper pools and the consequent disruption of cellular redox homeostasis through uncontrolled copper-mediated reactive oxygen species generation. Current therapies mainly depend on high-affinity copper chelation to lower metal burden which sometimes also strip copper from cuproproteins and may disturb physiological copper-dependent processes. It also does not directly suppress pathological copper reactivity i.e., free radical generation, a major driver of WD progression. To overcome these limitations, we have rationally designed Gua-Cu-3, a C₃-symmetric guanidinium-based molecule that can chelate labile copper without metal stripping from cuproproteins due to moderate binding affinity and it has intrinsic antioxidant activity within a single nanosheet-forming supramolecular self-assembly. Spectroscopic, calorimetric, and computational analyses revealed controlled multivalent copper coordination (K_d_ = 95.4 μM) while radical-scavenging and hydroxyl-radical inhibition assays further revealed potent redox-regulatory activity. In cellular models of copper overload, Gua-Cu-3 treatment reduces ATP7B trafficking from trans-Golgi network, confirming effective intracellular copper sequestration. This was accompanied by a marked reduction in oxidative stress, reduction of translocation of Nrf2 in nucleus and of HO-1 expression, thereby limiting lipid peroxidation which leads to restoration of cellular homeostasis, including improved lysosomal integrity, reduced endoplasmic reticulum stress, decreased mitochondrial superoxide levels, and diminished apoptosis. Importantly, the protective effects extended, beyond cultured cells, Gua-Cu-3 attenuates oxidative stress in ATP7B-homolog-deficient *C. elegans* and rescues copper-induced developmental defects in zebrafish, outperforming D-penicillamine, which is currently in use for Wilson disease management. These findings establish Gua-Cu-3 as a molecularly designed supramolecular copper-reactivity buffer that couples-controlled copper sequestration with redox regulation which is distinct from conventional copper depletion and provides a framework for treating Wilson disease and other disorders associated with metal dyshomeostasis and oxidative stress.

## Introduction

Wilson disease is an autosomal recessive disorder arising from loss-of-function mutations in the ATP7B gene, which impairs biliary copper export and drives progressive accumulation of labile copper within the liver and later in brain, and peripheral tissues^1^. The consequent rise in labile copper provokes oxidative stress, mitochondrial dysfunction, and hepatocellular injury, manifesting clinically as hepatic failure, neuropsychiatric impairment, movement disorders, and in some patients characteristic Kayser- Fleischer rings at the edge of the cornea^2^. Although, dysregulation of copper homeostasis leads to various diseases progression, copper is essential micronutrient which required for various physiological processes such as ATP production, iron homeostasis, antioxidant protection, and immune activity. Due to this dual functional nature, copper level is tightly regulated inside cells.^3^ The intracellular concentration of copper is kept at an optimum level through a buffering system to prevent toxicity, with copper primarily being present in a Cu(I) bound form by metallochaperones, glutathione, and metallothionein^4^. When this buffering capacity fails, as in Wilson disease redox active labile pool of copper takes part in the Cu(I)/Cu(II) redox cycle along with a free radical-mediated ligand-assisted Fenton-type reaction which further causes oxidative stress, leading to oxidative damage to proteins, lipids, and nucleic acid^5^. Thus, oxidative stress becomes an important amplifier of Wilson disease pathology. Uncontrolled oxidative stress disturbs cellular homeostasis by oxidizing protein, inducing endoplasmic stress, excessive autophagy and ultimately cell death^6^.

Current management mostly depends on copper chelators such as d-penicillamine, Trientine, etc., or zinc-based inhibition of absorption of copper via intestine^7^. While this has led to significant improvement in Wilson disease, it cannot reduce copper-mediated oxidative damage and is often linked with various side effects. d-penicillamine treatment sometimes leads to paradoxical neurological worsening, whereas Trientine therapy has led to some hemotoxicity in some cases and neurological regression^8^. Fundamentally, current treatments only aim to reduce the labile copper pool without regulating its pathological reactivity. As a result, the reduction of oxidative damage mediated by copper is only done indirectly, even though oxidative stress is an important part of copper pathologies. This shows that removing copper alone is not sufficient for managing its pathological impacts. New methods that target both labile copper availability and oxidative stress need to be found. Unfortunately, existing methods that try to do both, cannot give chemically controllable regulation of copper reactivity.

As an alternative, we hypothesized that a strategy that protects against pathological copper reactivity, without stripping copper from cuproproteins with antioxidant properties, could offer a solution. For such an alternative approach to work, it should involve a molecule that can specifically target and bind labile copper ions with moderate rather than strong affinity, while at the same time reducing oxidative stress. Notably, to achieve this goal, specific tuning of metal-binding and redox properties is necessary within a single molecular framework rather than conventional small-molecule chelators or repurposed antioxidants^9^.

Guided by this concept, we rationally designed Gua-Cu-3, a C3-symmetric guanidinium-based molecule that integrates three complementary functional elements within a single scaffold. First, ortho-hydroxylated imine motifs which provides multivalent copper-binding sites capable of controlled copper coordination. Second, redox-active phenolic groups which possess intrinsic antioxidant activity that can directly suppress reactive oxygen species. Third, the guanidinium-centered scaffold promotes supramolecular self-assembly into nanosheet structure, enabling multivalent presentation of copper-binding motifs while potentially enhancing serum stability. Together, these features were designed to transform Gua-Cu-3 from a conventional chelator into a molecularly designed copper-reactivity buffer capable of simultaneously regulating labile copper availability and oxidative stress.

Through spectroscopic, cellular, and *in vivo* investigations involving ATP7B-deficient cells, ATP7B-homolog-deficient *Caenorhabditis elegans*, and copper-overloaded zebrafish, we show that Gua-Cu-3 chelates intracellular copper, attenuates oxidative stress and its associated cellular dysfunction. It also alleviates copper-induced pathological phenotypes and most notably, rescues copper-induced developmental defect i.e., swim bladder inflation in zebrafish. To delineate the specific contribution of copper chelation, we synthesized a structurally analogous control molecule, Gua-3, which retains the guanidinium-centered scaffold and supramolecular assembly characteristics but lacks the ortho-phenolic hydroxyl groups required for efficient copper binding. Comparative studies between Gua-Cu-3 and Gua-3 enabled the decoupling of copper-chelation and antioxidant effects, thereby establishing the critical role of controlled copper coordination in mitigating copper-induced toxicity. Furthermore, the present study identifies copper reactivity buffering as an entirely new strategy, distinct from traditional copper chelation. It defines molecular design of multifunctional scaffolds as a valuable approach to treat Wilson disease and related disorders associated with metal-induced oxidative stress.

## Results

### Design-Strategy of Gua-Cu-3 and Gua-3

Although the concept of copper chelation is still relevant to Wilson disease (WD) treatment, clinically used chelators like D-Penicillamine (D-Pen) and Trientine exert their action based on the strong interaction with the exchangeable copper pool^12^. D-Pen produces highly stable complexes in the form of Cu(l)-thiolate (log β_504_ = 101.5 for the predominant species and log β_122_ = 39.18 for mononuclear complex), being much more stable than Cu(l)-glutathione complexes (log β = 38.8)^13^. Even though this property ensures high effectiveness, it can lead to the stripping of Cu⁺ from glutathione when D-Pen is used in high amounts (in the case of the d-pen dose being greater than 1 mM at later stages of Wilson disease). This can affect the buffering mechanism for intracellular copper and cuproproteins in general^14^. At the same time, it should be noted that the chelating action in itself cannot prevent copper-mediated oxidative damage, which occurs via Fenton-type reactions, which represents a major contributor to cellular injury in WD. This has led to the formulation of a two-pronged approach involving the controlled sequestration of a labile pool of copper ions along with built-in redox modulation capability. In our view, a single entity that can have moderate copper-binding properties along with built-in redox modulatory potential can help to control excessive levels of available labile copper ions, in addition to reducing any possible copper-induced oxidative stress. This concept may serve as an additional tool beyond the traditional method of chelation for dealing with both the metal disorder and its redox effect. Furthermore, embedding these functionalities within a self-assembling molecular framework could provide multivalent presentation of binding sites and enhance biological performance.

Accordingly, we designed a C₃-symmetric guanidinium-based scaffold, Gua-Cu-3 comprising three ortho-hydroxylated Schiff-base arms radiating from a central guanidinium core (Scheme 1a). The C₃-symmetric design was selected to maximize multivalent copper coordination while maintaining a compact molecular framework. The ortho-phenolic imine motifs were incorporated as copper-binding units, whereas the phenolic hydroxyl groups simultaneously serve as redox-active centers capable of scavenging reactive oxygen species. The permanently cationic guanidinium core was expected to facilitate cellular internalization and promote supramolecular self-assembly into two-dimensional nanosheet architectures. Collectively, The scaffold was engineered to exhibit moderate copper-binding affinity which is sufficient to target labile copper pools while minimizing the stripping copper from essential metalloenzymes while simultaneously attenuating copper-induced redox imbalance. To distinguish the contribution of copper coordination from the structural and supramolecular characteristics of the scaffold, Gua-3, which retains the guanidinium core and self-assembly behavior but lacks the ortho-phenolic hydroxyl groups required for effective copper binding. Comparative evaluation of Gua-Cu-3 and Gua-3 enables direct assessment of the therapeutic advantage conferred by the dual-function design in WD. Scheme 1a illustrates the structures of Gua-Cu-3 and the non-chelating control scaffold Gua-3, highlighting the presence or absence of the ortho-phenolic hydroxyl group responsible for copper coordination, as well as the anticipated oxidizable phenyl ring in Gua-Cu-3. Scheme 1b is a cartoon representation of 2D nano-sheets of Gua-Cu-3, exfoliation using ultrasound and binding to Cu (II)/Cu(I) ions in biological conditions. Subsequent experiments were undertaken to evaluate the physicochemical properties, copper-binding behavior, redox activity, and biological efficacy of these scaffolds in cellular and organismal models of Wilson disease.

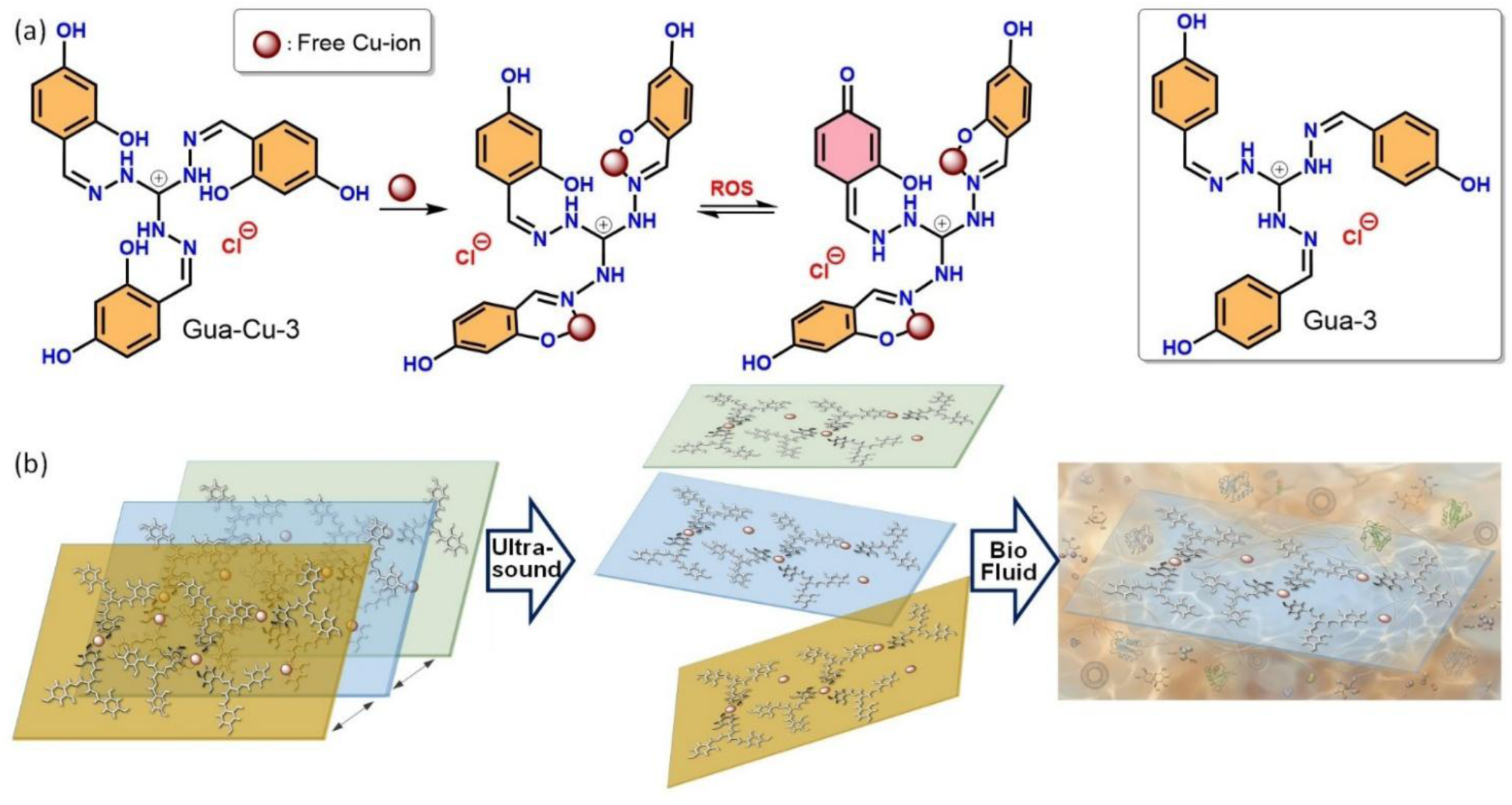

### Synthesis of Gua-Cu-3 and Gua-3

Guanidinium hydrazine was prepared following the published protocol^15^. Guanidinium hydrazine (1.0 equiv.) and 2,4-dihydroxybenzaldehyde (3.15 equiv.) were dissolved in EtOH/H₂O (95:5, v/v). The reaction mixture was refluxed overnight. Upon completion of the reaction, the solution was cooled to room temperature and stored at 4 °C for 36 h, during which a precipitate formed gradually. The solid was collected by filtration using Whatman 41 and washed several times with cold ethanol to remove unreacted starting materials. In parallel, a control compound, Gua-3, was synthesized which is structurally analogous but non-chelating in nature, and was employed as a structural control throughout the study. Gua-3 was synthesized under identical conditions using 4-hydroxybenzaldehyde in place of 2, 4-dihydroxybenzaldehyde. The two scaffolds were designed to differ primarily in the presence or absence of ortho-phenolic hydroxyl groups, enabling subsequent evaluation of the contribution of copper coordination to biological activity of the scaffold. Structural integrity and purity of both compounds were confirmed by HRMS and NMR spectroscopy (**Fig. S1-S5**).

### Structural characterization of Gua-Cu-3 and Gua-3 reveals their self-assembly into nanosheet architectures

To investigate the supramolecular behavior of the guanidinium-based scaffolds, Gua-Cu-3 and Gua-3 (1 mg each) were separately dispersed in 1 mL of 1X PBS buffer. The suspensions were sonicated for 45 min to ensure homogeneous dispersion and promote self-assembly. Following equilibration for 15 min at room temperature to allow equilibration, aliquots of the solutions were drop-cast onto carbon-coated copper TEM grids for transmission electron microscopy (TEM), silicon wafers for field-emission scanning electron microscopy (FE-SEM), and atomic force microscopy (AFM), and FTO-coated substrates for Kelvin probe force microscopy (KPFM). The samples were allowed to dry under ambient conditions before subsequent imaging. TEM, FE-SEM, and three-dimensional AFM images confirmed that both Gua-Cu-3 and Gua-3 spontaneously self-assemble into layered nanosheet-like architectures, which may facilitate interaction with biological environments. Furthermore, KPFM analysis revealed a positive surface potential for these assemblies, arising from the intrinsic cationic nature of the guanidinium scaffold (**Fig. 1 A-H**). PXRD analysis revealed distinct reflections at 2θ = 9.4°, 16.1°, 20.1°, and 26.6°, corresponding to d-spacings of 9.41, 5.51, 4.42, and 3.35 Å, respectively. The low-angle reflection at 9.4° is indicative of lamellar ordering, whereas the intense peak at 26.6° (d = 3.35 Å) is characteristic of aromatic π-π stacking. Together, these diffraction features support an ordered supramolecular nanosheet architecture stabilized through both lamellar organization and intermolecular aromatic interactions. (**Fig. 1I, Table-1**). The regular diffraction pattern supports the presence of well-defined supramolecular organization within the nanosheet assemblies. We found that during a 24-hour incubation period, despite the presence of fetal bovine serum, the Gua-Cu-3 compound still retained its colloidal stability, based on the observation that the supramolecular structure of Gua-Cu-3 remained intact (Fig. 1J, K; S6)^16^. Overall, the findings indicate that Gua-Cu-3 can form stable supramolecular structures suitable for further studies regarding the binding to copper ions.

**Figure 1.**
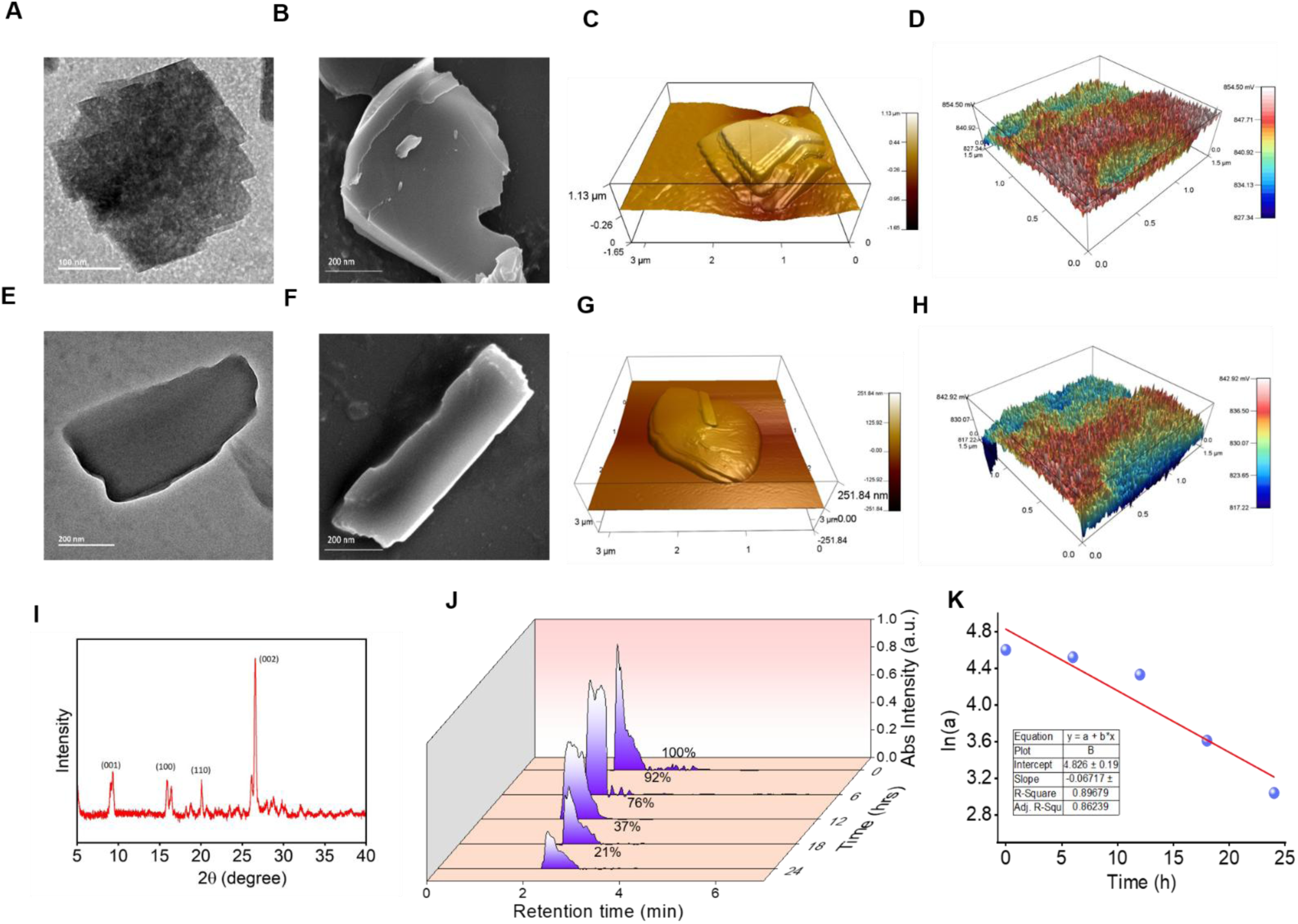
Structural characterization and serum stability of Gua-Cu-3 and Gua-3. (A-D) Morphological and surface potential characterization of Gua-Cu-3 assemblies. (A) TEM image showing well-defined layered nanosheet-like architectures. (B) FE-SEM image confirming sheet-like morphology with lateral dimensions in the nanometer regime. (C) Three-dimensional AFM topography image displaying stacked lamellar features. (D) KPFM surface potential map indicating a positive surface potential distribution across the nanosheets. (E-H) Corresponding characterization of Gua-3, demonstrating similar nanosheet-like self-assembly and positive surface charge distribution. (E) TEM and (F) FE-SEM images reveal similar layered nanosheet-like self-assembled structures. (G) Three-dimensional AFM topography image and (H) KPFM map further confirm lamellar morphology and a net positive surface potential, consistent with the cationic guanidinium scaffold. (I) PXRD profile of compound Gua-Cu-3 indicating the presence of characteristic diffraction peaks indicative of ordered lamellar structure. (J) Representative HPLC profiles indicating the stability of compound Gua-Cu-3 when kept in FBS for 24 hours. (K) Plot demonstrating kinetics of stability indicated by ln(a) vs time graph.

### Computational and Experimental Characterization of Copper Binding by Gua-Cu-3

To obtain an atomistic perspective on copper coordination, DFT was used to investigate the free ligand (Gua-Cu-3) and its corresponding Cu(II) complex. The optimized geometry showed that the interaction between the two is energetically favorable, as evident from the negative ΔE value (−17.65 kcal.mol^-1^), suggesting that a stable complex can be formed (**Fig. 2A & B**). This indicates that the metal ion is possibly coordinated through the ortho-phenolic oxygen and other coordinating sites in accordance with the suggested model. Importantly, the moderate binding energy obtained is ideal for the selective recognition and chelation of labile copper pools. Conceptual DFT calculations showed a significant electronic change upon copper binding. As compared to the ligand, there has been a substantial reduction in the value of the energy gap between HOMO and LUMO orbitals from 3.044 eV to 0.905 eV, along with a reduction in chemical hardness from 1.522 eV to 0.452 eV, and an increase in electrophilicity from 1.051 eV to 15.288 eV in the case of the copper-bound complex. Therefore, based on these values, it can be concluded that, with increased electronic delocalization and polarization driven by charge transfer, the formation of a coordination complex becomes more favorable (**Fig. S7A & B and Table 2**). Collectively, these calculations support the feasibility of copper binding by Gua-Cu-3 and provide a structural framework for subsequent experimental investigation.

**Figure 2.**
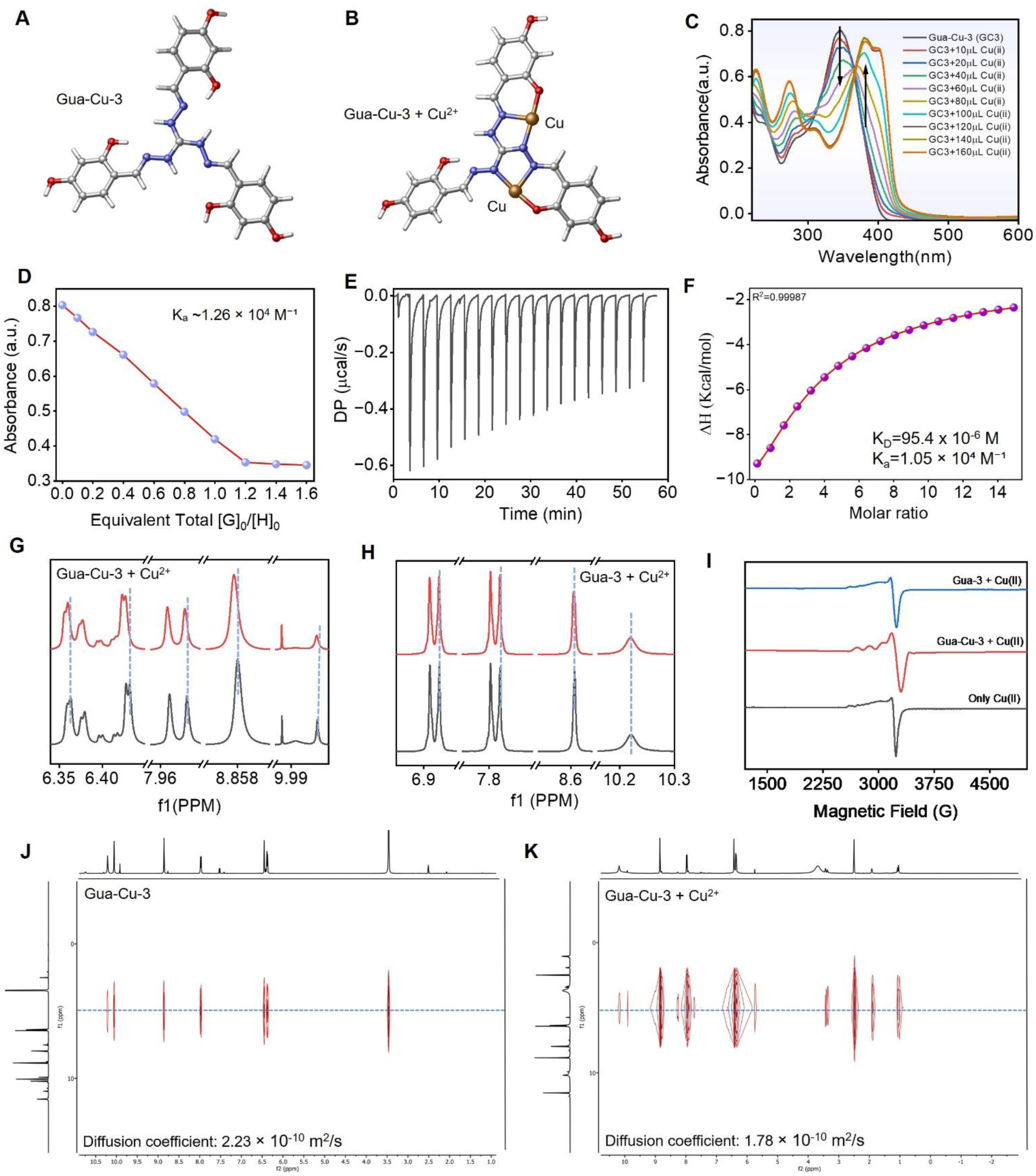
Computational and Experimental Characterization of Copper Binding by Gua-Cu-3. (A) GuaCu-3 optimized structure without Cu²⁺. (B) Coordination environment of Cu2+ within the GuaCu-3 complex, calculated using B3LYP/6-31G+(d,p) theory level. Colours represent the atoms as follows: nitrogen - blue, oxygen - red, copper - orange, and carbon - silver. (C) UV-vis titration results of Gua-Cu-3 in the presence of increasing concentration of Cu(II) leading to an evident decrease in the ∼350 nm absorbance band while simultaneously increasing the absorbance intensity of the red-shifted band centered at ∼400 nm with an isosbestic point of ∼385 nm, indicating efficient binding of copper ions. (D) Analysis of the UV-vis titration results by BindFit software indicating a defined interaction between Gua-Cu-3 complex and Cu(II). (E) Exothermic ITC thermograms of titration between Cu(II) and Gua-Cu-3 resulting in clearly identifiable peaks of heat release. (F) Heat integration and non-linear fitting of ITC results showing the interaction affinity and binding stoichiometry with n ≈ 2. (G) Partial 1H NMR spectrum of Gua-Cu-3 both without and with added Cu(II) showing perturbation of proton chemical shifts due to coordination. (H) Corresponding ^1^H NMR spectra of control Gua-3 showing minimal spectral changes upon Cu(II) addition. (I) EPR spectra of Cu(II) alone, Cu(II) with Gua-Cu-3, and Cu(II) with Gua-3, demonstrating the emergence of resolved hyperfine splitting only in the presence of Gua-Cu-3, consistent with formation of a defined inner-sphere copper coordination environment. (J, K) DOSY NMR indicates a difference in the diffusion coefficient of Gua-Cu-3 from 2.23×10^-10^ m^2^/s to 1.78×10^-10^ m^2^/s after Cu²⁺ complexation, which means they have an altered hydrodynamic radius. Together, these complementary analyses support direct Cu(II) coordination by Gua-Cu-3 and are consistent with formation of a defined copper-ligand complex that is not observed for the control scaffold Gua-3.

**Table 1.** PXRD peak positions, d-spacing values, relative intensities, and corresponding structural assignments for the self-assembled Gua-Cu-3 nanosheets.

| Peak Position (2 theta) | (d)-spacing (Å) | Relative Intensity | Most Likely (hkl) Assignment | Physical Structural Meaning |
| --- | --- | --- | --- | --- |
| (9.4°) | (9.41) | Medium | (001) | Lamellar periodicity arising from ordered molecular stacking |
| (16.1°) | (5.51) | Medium | (100) | In-plane lateral molecular packing |
| (20.1°) | (4.42) | Medium | (110) | Secondary in-plane packing interactions within the supramolecular lattice |
| (26.6°) | (3.35) | Very Strong (Max) | (002) | Aromatic π-π stacking between adjacent molecular layers |

**Table 2.** Key descriptors were calculated at B3LYP/6-31G +(d, p) level of theory summarized in table 2.

| Species | $E_H$ (eV) | $E_L$ (eV) | $\Delta E$ (eV) | $\mu$ (eV) | $\eta$ (eV) | $\omega$ (eV) |
| --- | --- | --- | --- | --- | --- | --- |
| Gua-Cu-3 | -4.052 | -1.008 | 3.044 | -2.530 | 1.522 | 1.051 |
| Gua-Cu-3+Cu<sup>2+</sup> | -5.713 | -4.808 | 0.905 | -5.2605 | 0.452 | 15.288 |

The interaction between Gua-Cu-3 and copper ions was first evaluated via the use of UV-vis titration, during the UV-vis titration of Gua-Cu-3 with increasing concentration. For Cu(II), systematic changes were observed with clear spectra corresponding to a defined binding event. This resulted in the disappearance of an absorption peak around 350 nm, while a new peak appeared around 400 nm, accompanied by a shoulder at about 410 nm, along with other peaks at about 260 and 200 nm. Importantly, an isosbestic point at 385 nm was observed in the titration curve, suggesting a clean conversion of the two main species, the unbound ligand (Gua-Cu-3) and bound copper complex Cu (II)-Gua-Cu-3 (**Fig. 2C**)^17^. The bathochromic shift in the UV spectrum during the process of copper coordination may be explained by the decrease in the energy gap between HOMO and LUMO levels, which confirms the presence of a defined interaction between Gua-Cu-3 and Cu(II). A quantitative evaluation of the titration data using the BindFitPlot model showed the existence of a defined interaction between Gua-Cu-3 and Cu(II) (**Fig. 2D**). To further verify the spectroscopic binding assay and quantitatively evaluate the interaction thermodynamics, ITC was performed.

Titration of Cu(II) into Gua-Cu-3 produced exothermic heat changes consistent with a specific binding event and a hallmark of stoichiometric complex formation (**Fig 2E**). Fitting of the binding isotherm yielded a binding constant (K_a_ = 1.05 × 10⁴ M⁻¹) and a favorable enthalpy change (ΔH= -48.7 kcal·mol⁻¹), indicating that copper coordination is enthalpy-driven under these conditions. Such a binding profile is consistent with the formation of stabilizing coordination bonds and favorable metal-donor interactions in the bound state. The fitted stoichiometry (n ≈ 2) suggested the possibility that each Gua-Cu-3 molecule can accommodate two copper ions (**Fig 2F**)^18^. Importantly, the measured affinity supports the intended design strategy of engaging labile copper without relying on the extremely high-affinity coordination characteristic of conventional copper chelators. As expected, no significant heat changes were observed, when titrate Gua-3 with Cu(II), indicating the absence of measurable copper binding (**Fig S7 C, D**). Site-specific copper coordination was further supported by ^1^H NMR spectroscopy, in which the addition of Cu(II) induced pronounced chemical shift perturbations for resonances proximal to the guanidinium-based chelating unit of Gua-Cu-3(**Fig 2G**). By comparison, an otherwise analogous control scaffold without the copper-binding moiety did not present any detectable spectral variations upon Cu(II) treatment, confirming that copper coordination is structurally encoded within the Gua-Cu-3 framework (**Fig 2H**). In concert with these observations, EPR spectroscopy uncovered significant changes in the Cu(II) spectral features on the addition of Gua-Cu-3, indicating changes in the coordination environment of copper. Both the electronic environment and ligand field, confirm direct coordination. As shown in **Fig. 2I**, free Cu(II) exhibited a broad, poorly resolved EPR signal characteristic of a hydrated copper (II) species in solution. Upon addition of Gua-Cu-3, this spectrum evolved into a well-resolved hyperfine-split spectrum, reflecting the formation of a rigid inner-sphere copper coordination environment^19^. Notably, no such hyperfine splitting was observed in the presence of Gua-3, confirming its inability to effectively bind Cu(II). To confirm complex formation, DOSY NMR experiments were done to observe changes in hydrodynamic radius upon copper binding. Diffusion coefficient of Gua-Cu-3 changed from 2.23* 10^-10^ m^2^/s (Free Gua-Cu-3) to 1.78 * 10^-10^ m^2^/s (Gua-Cu-3-copper complex) after Cu(II) addition (**Fig. 2J,2K**). The observed reduction in diffusion rate provides additional support for stable complex formation^20^. Taken together, the combined computational, spectroscopic, calorimetric, and magnetic resonance analyses provide convergent evidence that Gua-Cu-3 directly coordinates copper ions through the ortho-phenolic motif and adjacent donor sites. In contrast, the control scaffold Gua-3 lacks the structural elements required for effective copper coordination. These findings establish Gua-Cu-3 as a chemically defined copper-binding scaffold and provide the mechanistic basis for its subsequent evaluation in models of copper-induced oxidative stress.

### Gua-Cu-3 exhibits copper chelation in Wilson disease Cellular Models

We next examined whether these *in vitro* copper-binding properties translate into functional intracellular copper sequestration. First, we evaluated the safety profile of Gua-Cu-3 before testing its copper chelation property and therapeutic potential for Wilson disease. We utilized HepG2 cells lines (WT_HepG2), an established cellular model to study hepatic phenotypes. To create Wilson disease (WD) conditions, we used CRISPR-Cas9 generated ATP7B knock-out HepG2(ATP7B-/-_HepG2) using which serve as cellular model for Wilson disease^9^.

Gua-Cu-3 was found to be non-toxic up to 30 µM for at least 24 hours in both WT_HepG2 and ATP7B-/- _HepG2 cells (**Fig. S8A, S8B, S8C**) and for all biological experiment, the concentration did not exceed 25 µM. To evaluate intracellular copper chelation property, we have checked ATP7B sub-cellular localization. Cellular copper overload induces the trafficking of ATP7B from the TGN to vesicular compartments as part of the cellular copper-export response. In WT_HepG2 cells exposed to elevated extracellular copper (100 µM), treatment with Gua-Cu-3 largely preserved ATP7B localization at the TGN, indicating attenuation of copper-induced ATP7B redistribution and suggesting intracellular copper chelation (**Fig. 3A**).^22^ Quantitative analysis confirmed significant restoration of ATP7B colocalization with the TGN marker Golgin97. When compared with established copper chelators, Gua-Cu-3 demonstrated higher efficacy than tetrathiomolybdate (TTM) and comparable performance to bathocuproine disulfonate (BCS) (**Fig 3B**) in restoration of ATP7B localization at Golgi. This indicates robust copper sequestration within the cellular environment.

**Figure 3.**
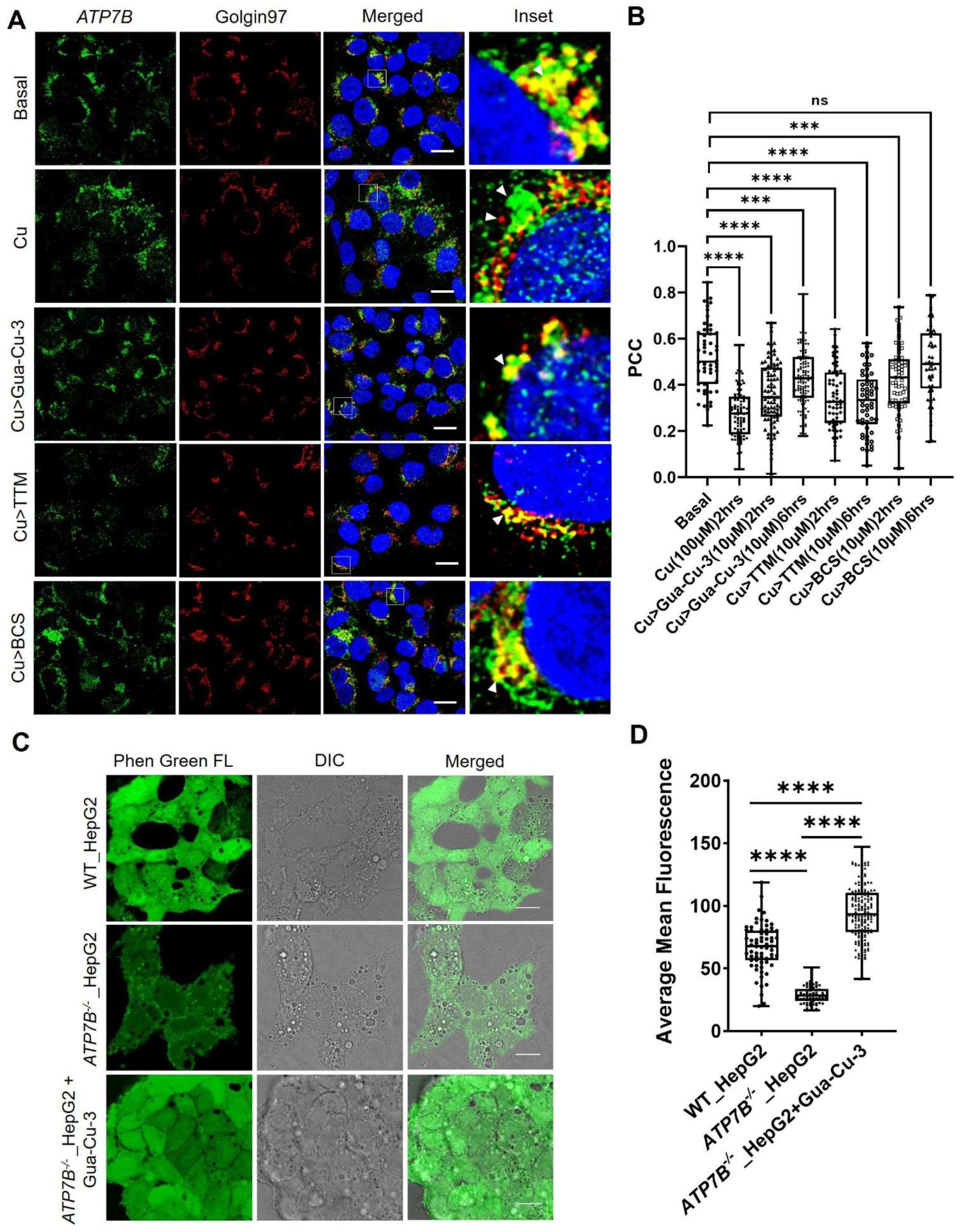
Gua-Cu-3 restores ATP7B localization and reduces labile copper in cellular models of Wilson disease. (A) Confocal fluorescence images of WT_HepG2 cells under basal conditions and following treatment with Cu (100 μM), Cu + Gua-Cu-3 (10 μM), Cu + tetrathiomolybdate (TTM, 10 μM), and Cu + bathocuproine disulfonate (BCS, 10 μM), immunostained for ATP7B (green), the trans-Golgi network marker Golgin97 (red), and nucleus (blue). Merged and magnified panels show ATP7B localization. Copper leads to the relocalization of ATP7B from the TGN to the endolysosmal and peripheral vesicles, while Gua-Cu-3 retains it in the TGN as a perinuclear structure, similar to reference chelators. Scale bars, 10 μm. (B) Colocalization of ATP7B and Golgin97 determined by PCC, shows the significant rescue of TGN localization by Gua-Cu-3 compared to copper treated, and equivalent or superior effect compared to the reference chelators. (C) Confocal microscopy images of intracellular labile copper levels in WT_HepG2, ATP7B-/-_HepG2, and ATP7B-/-_HepG2 exposed to Gua-Cu-3, labeled with the copper-sensing fluorophore Phen Green FL. DIC and overlay images were shown. Decreased fluorescence in ATP7B^-/-^_ HepG2 cells suggests high labile copper concentration, while Gua-Cu-3 rescues the fluorescence in ATP7B^-/-^_ HepG2 cells. Scale bars, 20 μm. (D) Intensity of the fluorescence measured using Phen Green is rescued by Gua-Cu-3 in ATP7B⁻/⁻_ HepG2 cells, suggesting a reduction in the level of labile copper. In conclusion, Gua-Cu-3 is efficient in the reduction of intracellular labile copper and restores ATP7B localization. Box plots represent medians along with their 25th to 75th percentiles, while whiskers represent the data points up to 1.5 times the the interquartile range. The non-parametric tests (Mann-Whitney U-test) were performed for comparing unpaired groups. Statistical significance is defined by *p<0.05, **p<0.01, ***p<0.001, ****p<0.0001, while ns stands for not significant. Statistical analysis and graphs were done using GraphPad prism software. Code for ImageJ macros can be found at GitHub.

To further assess changes in labile intracellular copper upon Gua-Cu-3 treatment in cellular model of Wilson disease i.e., ATP7B deficient HepG2(ATP7B-/-_HepG2), we employed the copper- sensitive fluorophore Phen Green. Comparing to WT_HepG2 cells, copper levels remain high in ATP7B-deficient HepG2 cells as reported earlier and henceforth copper accumulation quenched Phen Green fluorescence in ATP7B-/- _HepG2 cells, whereas treatment with Gua-Cu-3 restored Phen Green fluorescence intensity, indicating effective intracellular copper sequestration (**Fig. 3C,3D**)^23^. As a whole, these assays indicate the functional relevance of Gua-Cu-3’s copper-binding capabilities by showing that it increases ATP7B colocalization with TGN and restores Phen Green fluorescence, suggesting that it can interact with excess labile copper within cells in a Wilson disease-like environment.

### Gua-Cu-3 Suppresses Free Radical Generation and Copper-Driven Oxidative Chemistry

Having established that Gua-Cu-3 chelates copper both *in vitro* and *in cellulo*, we next investigated whether this interaction can be translated into the suppression of copper-driven oxidative chemistry. The antioxidant capacity of Gua-Cu-3 was assayed using DPPH radical scavenging activity, in which Gua-Cu-3 showed a concentration-dependent decrease in DPPH absorbance demonstrating efficient free radical scavenging activity comparable to the reference antioxidant ascorbic acid (**Fig. 4A**)^24^. By contrast, the structurally related control scaffold Gua-3, which lacks a copper-binding function also exhibited radical scavenging ability but less efficient compare to Gua-Cu-3. This result was further confirmed by the ABTS•⁺ assay, wherein Gua-Cu-3 was an effective quencher of the radical cation at all concentrations tested, and Gua-3 exhibited comparatively lower quenching activity (**Fig. 4B**)^25^. Collectively, these assays establish that Gua-Cu-3 possesses intrinsic antioxidant properties arising from its chemical structure.

**Figure 4.**
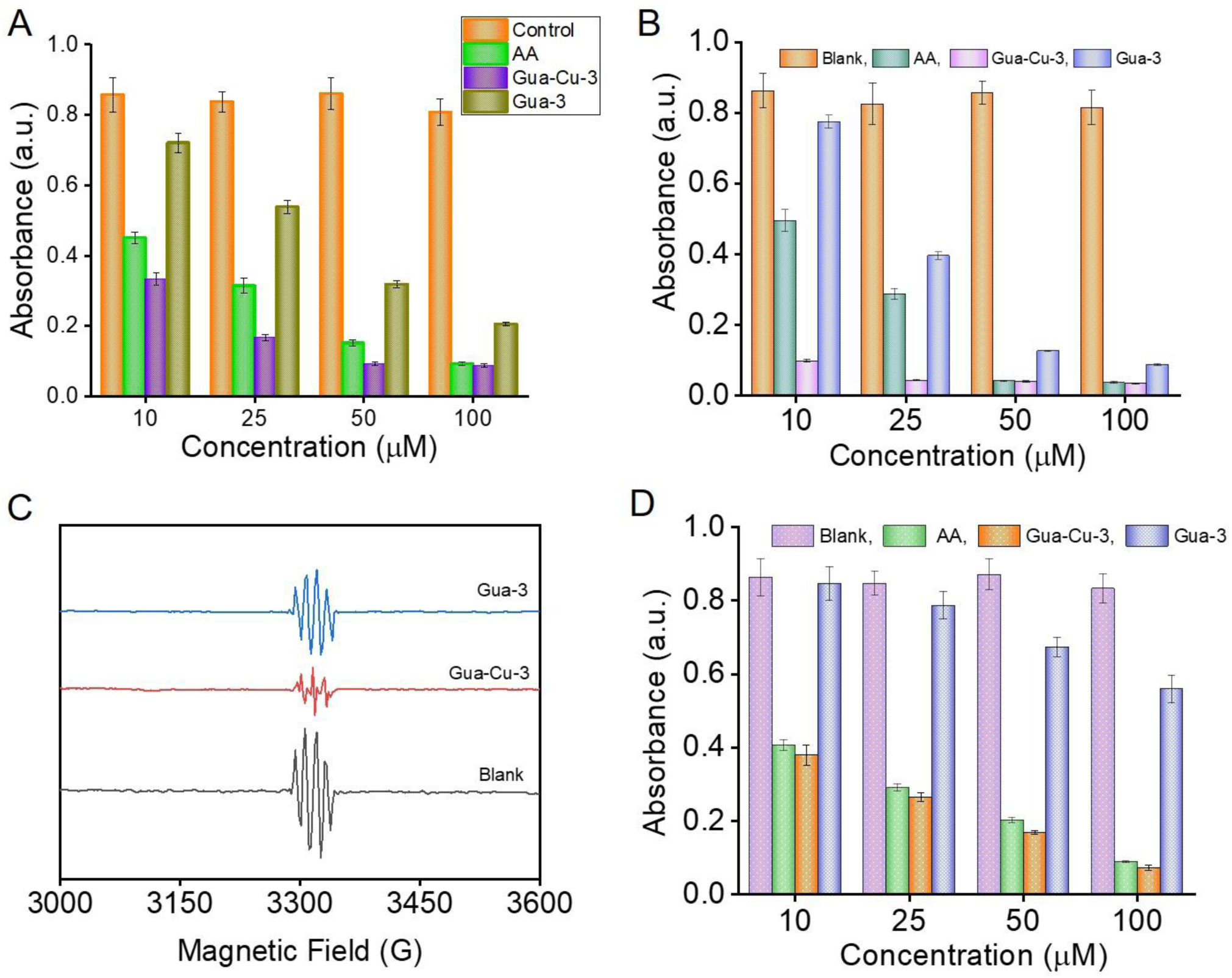
Gua-Cu-3 suppresses free radical generation and copper-driven oxidative chemistry. (A) DPPH radical scavenging assay displaying absorbance decrease following exposure to Gua-Cu-3, in a concentration-dependent manner similar to that observed for the reference antioxidant, ascorbic acid (AA); however, Gua-3 showed minimal scavenging activity at lower concentration. Lower absorbance levels denote higher radical scavenging efficiency. (B) Quenching of the ABTS•⁺ radical cation decolorization reaction in which Gua-Cu-3 shows considerable scavenging activity at all concentrations tested, equivalent to that seen with AA but greater than that observed for Gua-3, thus validating intrinsic antioxidant potential. (C) EPR spectrum showing Gua-Cu-3-mediated attenuation of the DMPO spin-trapped hydroxyl radicals (DMPO-OH) under copper-mediated Fenton-type reaction conditions (in absence/presence of Gua-Cu-3). Gua-3 does not show much significant effect, which demonstrates selective quenching of the radicals generated under Cu(II)-mediated oxidative stress. (D) Deoxyribose degradation assay to measure hydroxyl radical-induced oxidation of 2-deoxy-D-ribose. Inhibitory effect against oxidation, as shown through decreased absorbance of thiobarbituric acid-reactive species (TBARS) formation, was observed only for Gua-Cu-3. These assays provide evidence that Gua-Cu-3 is an inherent antioxidant capable of suppressing the formation of hydroxyl radicals by copper ions.The enhanced activity of Gua-Cu-3 relative to the non-chelating control scaffold supports a cooperative contribution of copper coordination and radical-scavenging properties to its overall redox-modulating function.

While DPPH and ABTS assays assess direct radical-scavenging activity, they do not directly evaluate suppression of copper-dependent ROS generation. Therefore, we next investigated the ability of Gua-Cu-3 to quench highly reactive oxygen species generated by metal-dependent pathways using the deoxyribose degradation assay and EPR spectroscopy with DMPO as a spin-trapping agent. Under copper-driven ROS-generating conditions, a characteristic DMPO-OH signal was readily observed. Addition of Gua-Cu-3 strongly reduces the typical DMPO-OH peak. These results suggest that Gua-Cu-3 is able to effectively suppress hydroxyl radical formation (**Fig. 4C**)^26^. In contrast, Gua-3 produced minimal changes in the EPR spectrum, demonstrating that antioxidant activity alone is insufficient to effectively inhibit copper-mediated ROS generation.

These findings suggest that coupling antioxidant activity with copper-binding capability provides a greater capacity to suppress copper-dependent ROS generation than antioxidant activity alone. To determine whether suppression of hydroxyl radical formation translates into protection against oxidative biomolecular damage, we employed deoxyribose degradation assay as a functional measure of hydroxyl radical-mediated oxidation of a DNA-relevant substrate. In this system, copper-driven Fenton-type chemistry generates •OH, which attacks 2-deoxy-D-ribose (a surrogate for the sugar backbone of DNA). Oxidative fragmentation of deoxyribose produces reactive aldehyde-containing products (classically including malondialdehyde-like species) that can be derivatized with thiobarbituric acid (TBA) to form a chromogenic adduct quantified spectrophotometrically. Thus, higher absorbance reflects greater •OH-dependent sugar damage. Under copper/ROS-generating conditions, robust deoxyribose degradation was observed. Addition of Gua-Cu-3 suppressed this signal in a dose-dependent manner, indicating decreased hydroxyl radical-mediated sugar oxidation^27^. In contrast, Gua-3 failed to reduce deoxyribose degradation, supporting the conclusion that inhibition is not due to non-specific antioxidant effects of the scaffold (**Fig. 4D**). This experiment confirms the presence of two cooperative roles within a single molecular entity in Gua-Cu-3: redox capabilities and selective copper coordination. In comparative studies with Gua-3, it becomes evident that an inherent antioxidant role alone is insufficient to inhibit copper-mediated oxidative damage effectively. On the contrary, the better behavior observed in Gua-Cu-3 arises from a synergy between the copper sequestration role and redox modulation, making this approach biologically significant.

### Gua-Cu-3 Restores Cellular Viability by Suppressing Copper-Induced Oxidative Damage Observed in Wilson disease

Copper accumulation in Wilson disease patients’ liver triggers cell death, with oxidative stress and lipid peroxidation being major contributors^28,29^. Moreover, conventional chelation therapy may transiently mobilize redox-active copper pools, potentially exacerbating oxidative stress-associated pathology^30,31^. As Gua-Cu-3 has the ability to chelate copper and suppress copper-driven radical formation, we next tested whether these properties translate into protection against copper-induced cytotoxicity in a cellular model of Wilson disease. Exposure of HepG2 cells to elevated copper concentrations resulted in a pronounced reduction in cell viability, as evaluated by MTT assay, consistent with copper-induced cellular damage. Treatment with Gua-Cu-3 significantly rescued cell viability indicating that cytoprotection arises from the functional properties of Gua-Cu-3 (**Fig. 5A**). To directly assess whether intracellular oxidative stress is the major driver of copper induced cell death and improvement is occurring due to reduction in oxidative stress, we employed CellROX green staining followed by confocal microscopy and flow cytometric analysis. Copper overload led to a marked increase in CellROX fluorescence, reflecting elevated reactive oxygen species in both WT_HepG2 cells as well as ATP7B-/-_HepG2 cells. Notably, Gua-Cu-3 treatment reduced oxidative stress to near-basal levels whereas Gua-3 produced only a partial reduction in oxidative stress, suggesting that effective attenuation of oxidative stress requires the combined contributions of copper modulation and radical-scavenging activity (**Fig. 5B-E**)^32^. Reduction in oxidative stress by Gua-Cu-3 was further confirmed quantitatively by flow cytometry (**Fig. 5F,5G**). As lipid membranes represent a primary downstream target of copper-mediated oxidative injury, we next examined lipid peroxidation. Copper exposure induced robust lipid peroxidation, whereas treatment with Gua-Cu-3 significantly inhibited this process, consistent with attenuation of Fenton-type chemistry and secondary radical propagation (**Fig. 5H**)^33^. The ability of Gua-Cu-3 to simultaneously reduce oxidative stress and lipid peroxidation provides a mechanistic basis for the observed recovery in cell viability.

**Figure 5.**
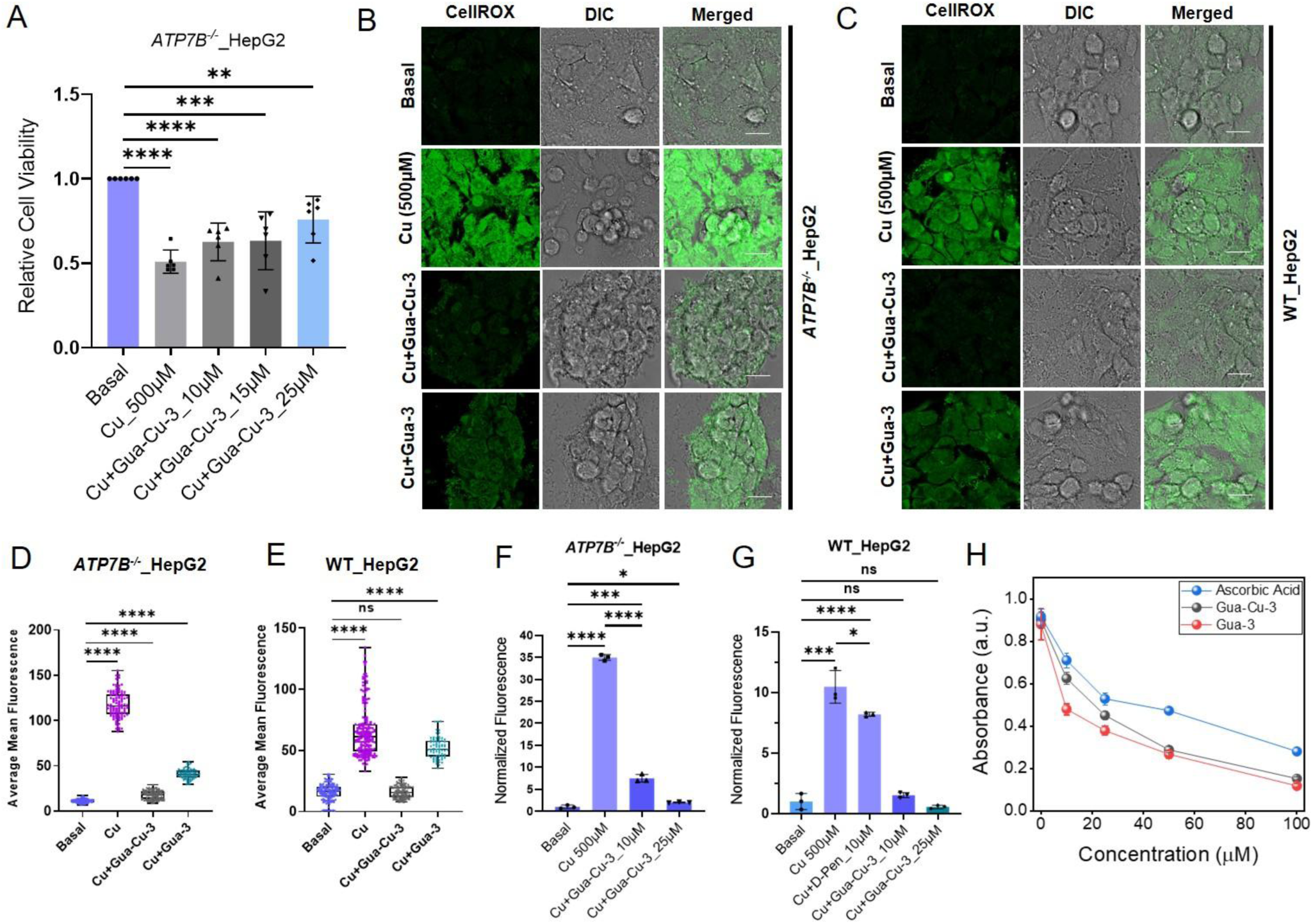
Gua-Cu-3 protects against copper-induced oxidative stress, lipid peroxidation. (A) Cell viability of ATP7B-/-_HepG2 cells following exposure to Cu (500 μM) in the absence or presence of increasing concentrations of Gua-Cu-3, measured by MTT assay. Gua-Cu-3 significantly improves cell viability in a dose-dependent manner. (B) Representative confocal microscopy images of CellROX Green staining of ATP7B-/-_HepG2 cells under basal conditions and following treatment with Cu (500 μM), Cu + Gua-Cu-3, or Cu + Gua-3. Copper overload induces strong ROS accumulation, whereas Gua-Cu-3 markedly reduces oxidative stress. Corresponding DIC and merged images are shown. (C) CellROX Green confocal images of WT_HepG2 cells under the mentioned treatment conditions, shows suppression of copper-induced ROS by Gua-Cu-3 but not by Gua-3. (D) Quantification of CellROX fluorescence intensity in ATP7B-/-_HepG2 cells, confirming significant reduction in intracellular ROS levels upon Gua-Cu-3 treatment compared to copper alone. (E) Quantification of CellROX fluorescence intensity in WT_HepG2 cells, demonstrating effective ROS suppression by Gua-Cu-3. (F) Flow cytometry analysis of ROS levels in ATP7B-/-_HepG2 cells under copper overload, confirming dose-dependent reduction of oxidative stress by Gua-Cu-3. (G) Flow cytometry quantification of ROS levels in WT_HepG2 cells treated with Cu in the presence of D-penicillamine or Gua-Cu-3, demonstrating comparable ROS suppression by Gua-Cu-3. (H) Lipid peroxidation analysis showing that Gua-Cu-3 significantly inhibits copper-induced membrane oxidative damage compared to copper-treated controls. Collectively, these results demonstrate that Gua-Cu-3 protects cells from copper-induced oxidative injury by suppressing intracellular ROS accumulation, inhibiting lipid peroxidation, and restoring cellular viability, consistent with its dual copper-chelating and redox-modulating functions. Results are expressed as the mean ± S.D. obtained from three or more independent experiments. Statistical significance was determined by means of unpaired t-tests. *p < 0.05; **p < 0.01; ***p < 0.001.

Collectively, these findings validate that Gua-Cu-3 protects cells from copper-induced cytotoxicity by functionally decoupling copper accumulation from oxidative damage. Importantly, the limited efficacy of Gua-3 across all assays confirms that cytoprotection is not a generic property of the scaffold but instead arises from the integrated copper-chelating and redox-modulating features of Gua-Cu-3. These results establish a mechanistic link between labile copper modulation, oxidative stress suppression, and restoration of cellular viability under conditions relevant to Wilson disease.

### Gua-Cu-3 alleviates Stress-Response Pathways in Cellular Models of Wilson disease

Chronic copper overload is known to perturb multiple cellular stress-response pathways, including redox signaling, lysosomal homeostasis, mitochondrial redox balance and endoplasmic reticulum (ER) integrity^34,35,36^. Having established that Gua-Cu-3 chelates copper, suppresses copper dependent oxidative chemistry, and restores viability, we next examined whether these effects extend to broader markers of cellular health in both wild-type and ATP7B^⁻/⁻^ _HepG2 cells. Nuclear factor erythroid 2-related factor 2 (Nrf2) is a master transcription factor and central regulator of the cellular antioxidant response that translocates from the cytoplasm into the nucleus in response to oxidative stress. Once inside, Nrf2 binds to antioxidant response elements (ARE), initiating the transcription of antioxidant genes^37^. Consistent with elevated ROS levels, copper exposure resulted in increased nuclear accumulation of the transcription factor Nrf2 in both WT_HepG2 and ATP7B^⁻/⁻^ _HepG2 cells, as expected with oxidative stress signaling. Treatment with Gua-Cu-3 significantly reduced Nrf2 nuclear localization under conditions of copper overload, and restoring its distribution to basal levels (**Fig. 6A**). Quantitative analysis confirmed a significant reduction in nuclear Nrf2 levels in both WT and ATP7B^⁻/⁻^ _HepG2 cells, indicating the efficient suppression of the copper oxidative stress pathway along with free radical quenching (**Fig. 6B,6C**).

**Fig 6.**
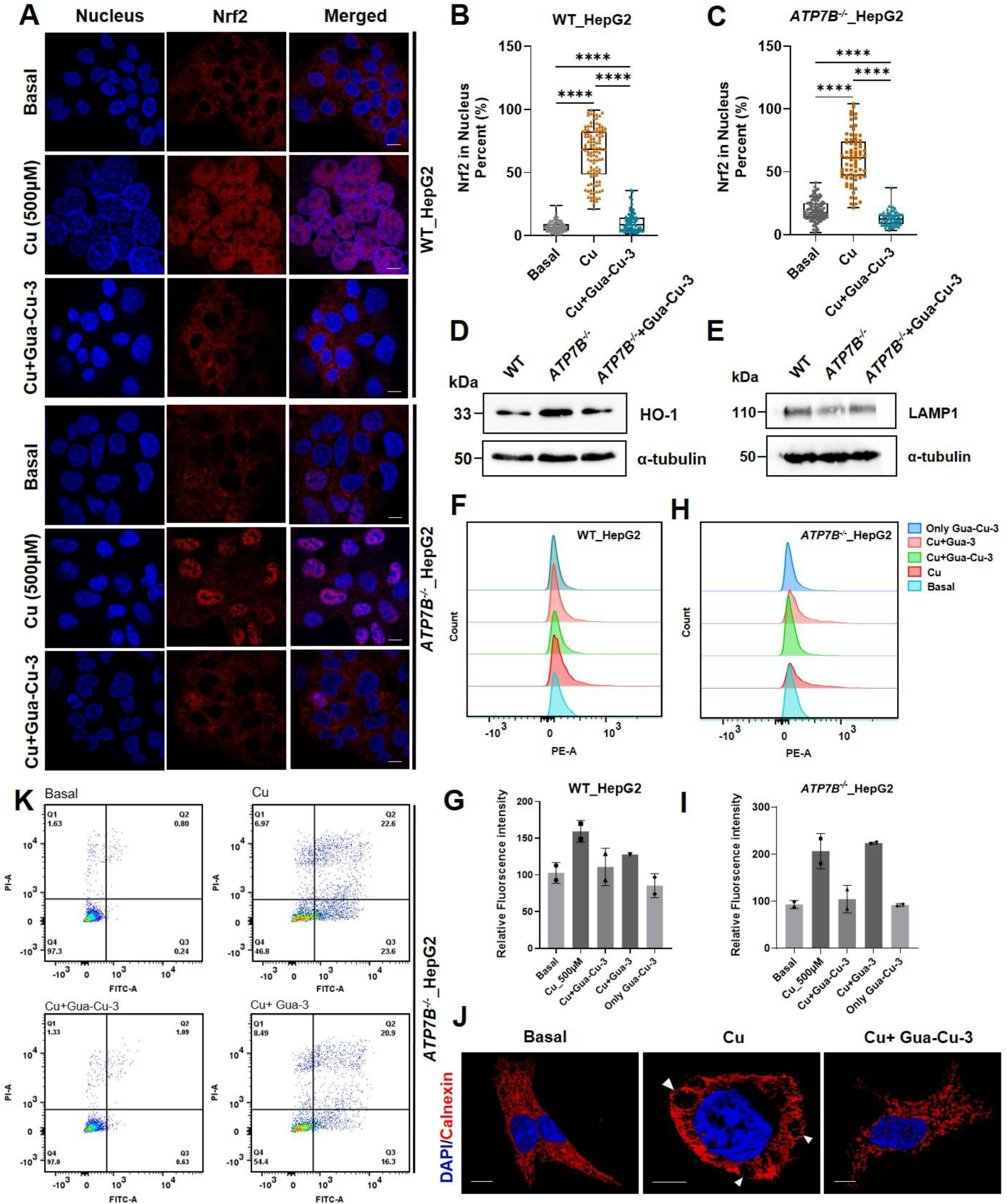
Gua-Cu-3 restores cellular stress-response pathways and organelle homeostasis under copper overload. (A) Confocal microscope images representative for Nrf2 distribution (Red) and nuclei (Blue) in WT and ATP7B⁻/⁻_HepG2 cells in basal conditions as well as after Cu (500 μM) treatment with or without Gua-Cu-3. Copper exposure causes Nrf2 translocation to the nucleus, while Gua-Cu-3 results in a re-localization of Nrf2 predominantly into the cytoplasm. Scale bar: 10 μm. (B, C) Nrf2 nuclear content in WT_HepG2 (B) and ATP7B⁻/⁻_HepG2 (C) cells, quantified as the ratio (%) to total Nrf2 protein, confirming significant reduction following Gua-Cu-3 treatment. (D) Representative Immunoblot image of the Nrf2 downstream target heme oxygenase-1 (HO-1) in WT, ATP7B-/-_HepG2, and ATP7B⁻/⁻_HepG2 cells treated with Gua-Cu-3, with α-tubulin as a loading control, showing normalization of stress-induced HO-1 expression. (E) Representative Immunoblot image of the lysosomal marker LAMP1 in WT, ATP7B^⁻/⁻^, and ATP7B^⁻/⁻^_HepG2 cells treated with Gua-Cu-3, demonstrating restoration of lysosomal protein levels. (F, H) Histograms representative for flow cytometric analysis of mitochondria superoxide production quantified with the aid of MitoSOX fluorescence in WT and ATP7B knockout HepG2 cells. (G, I) Quantification of mitochondria superoxide production determined using MitoSOX fluorescence in ATP7B knockout and WT_HepG2 cells, indicating significant inhibition of copper-induced mitochondrial ROS production by Gua-Cu-3. (J) Confocal fluorescence images of endoplasmic reticulum morphology visualized by calnexin staining (red) and nuclei (blue) in HepG2 cells, showing copper-induced ER structural disruption and whorl formation that is restored by Gua-Cu-3. Scale bars, 10 μm. (K) Flow cytometry analysis of apoptosis (Annexin V/PI staining) in ATP7B^⁻/⁻^ HepG2 cells under basal, copper-treated, and Gua-Cu-3 rescue conditions, demonstrating reduced apoptotic_HepG2 cell populations following Gua-Cu-3 treatment. Together, these data show that Gua-Cu-3 alleviates copper-induced cellular stress by suppressing Nrf2 activation, restoring lysosomal and mitochondrial function, preserving ER integrity, and preventing apoptosis, consistent with upstream attenuation of copper-driven oxidative damage. Results are expressed as the mean ± S.D. obtained from three or more independent experiments. Statistical significance was determined by means of unpaired t-tests. *p < 0.05; **p < 0.01; ***p < 0.001.

In agreement with Nrf2 localization data, expression of the downstream stress-responsive protein, heme oxygenase-1 (HO-1), was reduced following Gua-Cu-3 treatment (**Fig. 6D, S9A**)^38^. The coordinated reduction in Nrf2 activation and HO-1 expression supports the conclusion that Gua-Cu-3 lowers the oxidative burden associated with copper overload. Because oxidative stress is known to disrupt organellar homeostasis, we next examined the status of lysosomes, mitochondria, and the ER. Copper overload disrupted lysosomal integrity, indicated by a reduction in the levels of LAMP-1^39^; however, this was normalized by Gua-Cu-3 treatment (**Fig. 6E, S9B**). As mitochondria are the major source of free radical generation under metal toxicity and mitochondrial superoxide is the major contributor, next, we assessed mitochondrial superoxide using MitoSOX dye and observed a sharp increase in mitochondrial superoxide upon copper treatment which was attenuated upon Gua-Cu-3 treatment^40^ in both WT_HepG2 and ATP7B^−/−^_HepG2 cells (**Fig. 6F-I**). We next assessed ER stress by examining the formation of ER whorls, concentric membrane structures that arise under conditions of sustained oxidative and proteotoxic stress.^41^ Copper overload induced prominent ER whorl formation, indicative of ER dysfunction which was greatly reduced upon Gua-Cu-3 treatment (**Fig. 6J**). Given the widespread disruption of multiple organellar systems under copper overload conditions, we finally examined whether these cellular stresses culminated in apoptotic cell death. ATP7B-/-_HepG2 cells displayed a substantial increase in the apoptotic population following copper exposure. Treatment with Gua-Cu-3 significantly reduced apoptosis, consistent with its ability to alleviate upstream oxidative and organellar stress (**Fig. 6K**)^42^.

Overall, the results suggest that the protective activity of Gua-Cu-3 extends beyond lowering labile copper levels within cells. Through its antioxidant properties, Gua-Cu-3 regulates several stress response pathways, restores the normal functioning of various organelles, such as lysosomes, ER, mitochondria, and prevents apoptosis. The normalization of redox signals, lysosomal integrity, mitochondrial function, and endoplasmic reticulum morphology suggests that the regulation of oxidative chemistry by Gua-Cu-3 may play a key role in the pathogenesis of Wilson disease.

### Gua-Cu-3 alleviates copper-mediated oxidative stress and developmental defects across in small organism model

In order to find out whether the chelation capabilities and redox regulatory functions of Gua-Cu-3 have more extensive applicability outside the cellular model, we used complementary animal models for testing the ability of Gua-Cu-3 to correct the imbalance of copper *in vivo* systems, including both invertebrates and vertebrates. Initially, we tested the compound against the *C. elegans* mutant strain cua-1(tm12763), which lacks the homolog of human ATP7B and has copper content and oxidative stress^43,44^. The level of reactive oxygen species was measured via the CellROX staining. The tm12763 strain was shown to exhibit higher fluorescence signals compared to control worms. Gua-Cu-3 (10 µM, 12 hours) efficiently reduced the amount of detected ROS (Fig. 7A, 7B). To test whether the effects of Gua-Cu-3 in correcting copper-induced oxidative stress could be observed in vertebrate animals, we proceeded with a zebrafish (*Danio rerio*) embryo copper toxicity model^45,46,47^. Previous studies have demonstrated that excess copper disrupts swim bladder development, in part through mechanism linked to oxidative stress and impaired development signalling pathways, providing a sensitive *in vivo* phenotypic readout of copper toxicity^48^. Copper treatment produced the expected non-inflated swim bladder phenotype (**Fig. 7C).** To evaluate the prophylactic potential of Gua-Cu-3, 2 days post-fertilization (dpf) embryos were pre-treated with Gua-Cu-3 (0.5 µM) for 24 h, followed by copper exposure for 48 h. Notably, Gua-Cu-3 pre-treatment exhibited substantial restoration of normal swim bladder development (**Fig. 7D,7E**). Extensive washing prior to copper exposure minimized the possibility of residual compound effects, confirming that the observed rescue arises from a genuine protective mechanism. To directly link phenotypic rescue with redox modulation, we employed an acute oxidative stress model. Zebrafish embryos (3 dpf) were exposed to 200 µM copper for 45 min, resulting in a ∼2-fold increase in ROS levels as measured by CellROX Orange^49,50^. Pre-treatment with Gua-Cu-3 (10 µM, 30 min) effectively suppressed ROS accumulation to near-basal levels. Notably, under the experimental conditions employed, Gua-Cu-3 produced greater suppression of copper-induced ROS generation than D-penicillamine. (**Fig. 7F, 7G**).

**Figure 7.**
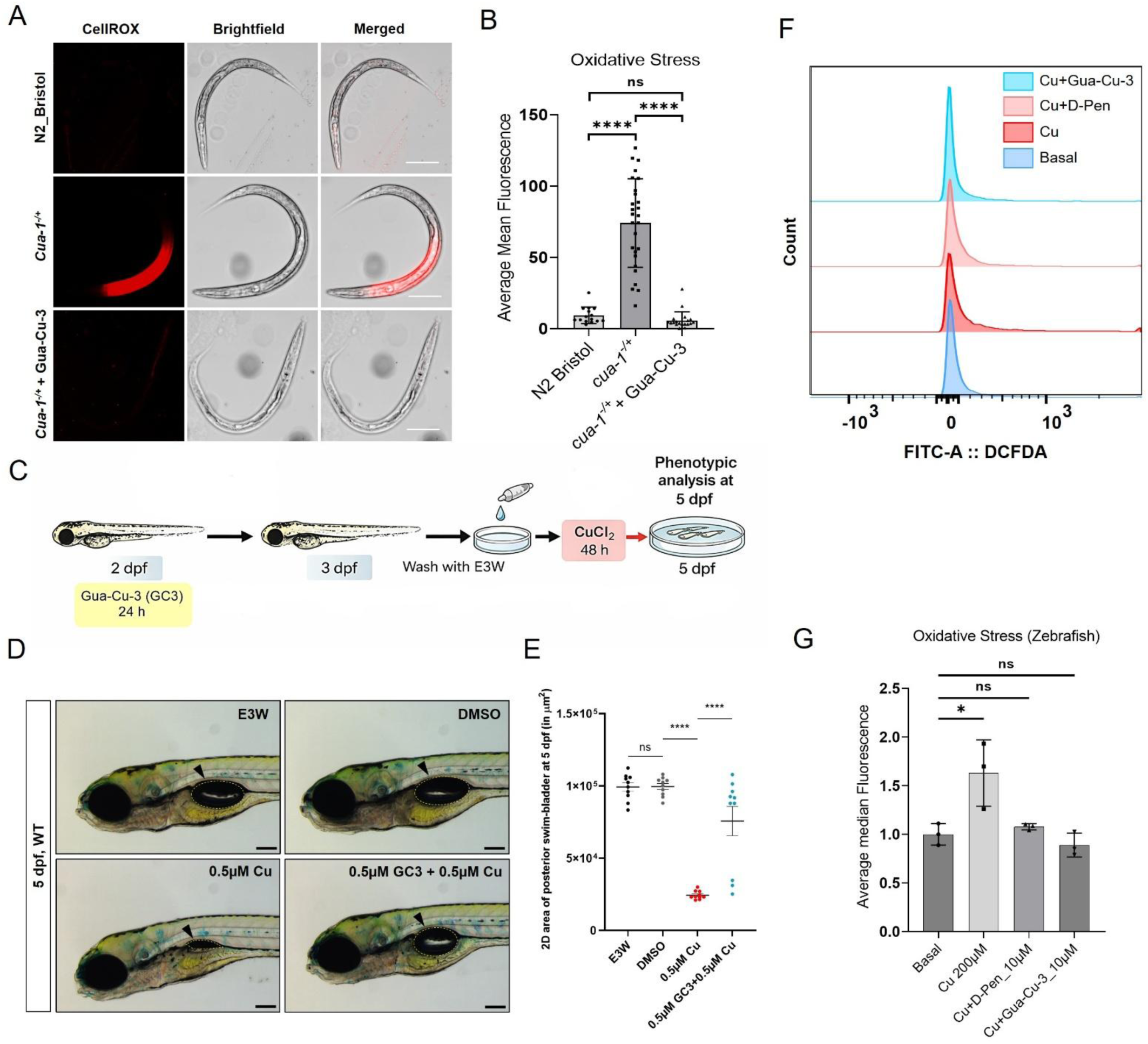
Gua-Cu-3 alleviates copper-induced oxidative stress and developmental defects across *in vivo* models. (A) Representative fluorescence images of CellROX staining in wild-type (N2 Bristol), copper-accumulating mutant (cua-1⁻/⁺, tm12763), and Gua-Cu-3-treated cua-1⁻/⁺ *C. elegans*, showing elevated oxidative stress in mutant worms and restoration toward basal levels following Gua-Cu-3 treatment. Corresponding brightfield and merged images are shown. Scale bars, 200 μm. (B) Quantification of oxidative stress levels in wild-type, cua-1⁻/⁺, and Gua-Cu-3-treated cua-1⁻/⁺ worms, showing significant reduction in ROS upon Gua-Cu-3 treatment. Data represent mean ± SD; statistical significance indicated as shown. (C) Schematic of Experimental schematic for copper and Gua-Cu-3 treatments in zebrafish embryos. (D) Brightfield images showing lateral views of 5 dpf larvae that were treated with copper and/or pre-treated with Gua-Cu-3 (the area of posterior swim-bladder was demarcated with dotted yellow line) (E) Quantification of swim bladder size in 5 dpf larvae (n=10). Data have represented as mean ± S.E.M. Statistical analysis by one-way ANOVA post-hoc Tukey’s test, P<0.0001****; Scale bar: 200µm. (F) Representative flow cytometry histograms of cellRox orange fluorescence in Zebrafish populations under the indicated conditions, demonstrating increased ROS in Zebrafishand suppression following Gua-Cu-3 treatment. (G) Quantification of oxidative stress in zebrafish embryos (3 dpf) measured using CellROX Orange fluorescence following exposure to Cu (200 μM) in the absence or presence of Gua-Cu-3 (10 μM) or D-penicillamine (D-Pen, 10 μM) (n= 10). Copper exposure significantly increases ROS levels, whereas Gua-Cu-3 reduces oxidative stress to near-basal levels and shows comparable or improved efficacy relative to D-Pen. Together, these results demonstrate that Gua-Cu-3 effectively alleviates copper-induced oxidative stress in both invertebrate and vertebrate models, confirming its protective activity across organismal systems. Results are expressed as the mean ± S.D. obtained from three or more independent experiments. Statistical significance was determined by means of unpaired t-tests. *p < 0.05; **p < 0.01; ***p < 0.001.

Collectively, these findings demonstrate that the protective effects of Gua-Cu-3 are maintained across diverse biological systems ranging from genetically defined invertebrate models to vertebrate organisms. The consistent reduction in oxidative stress together with rescue of copper-induced developmental abnormalities supports a mechanism in which modulation of copper-associated toxicity and oxidative damage contributes to organismal protection. When considered alongside the cellular and biochemical data, these results establish Gua-Cu-3 as a multifunctional scaffold capable of attenuating copper-induced oxidative stress across multiple levels of biological complexity **(Fig. 8).**

**Fig. 8.**
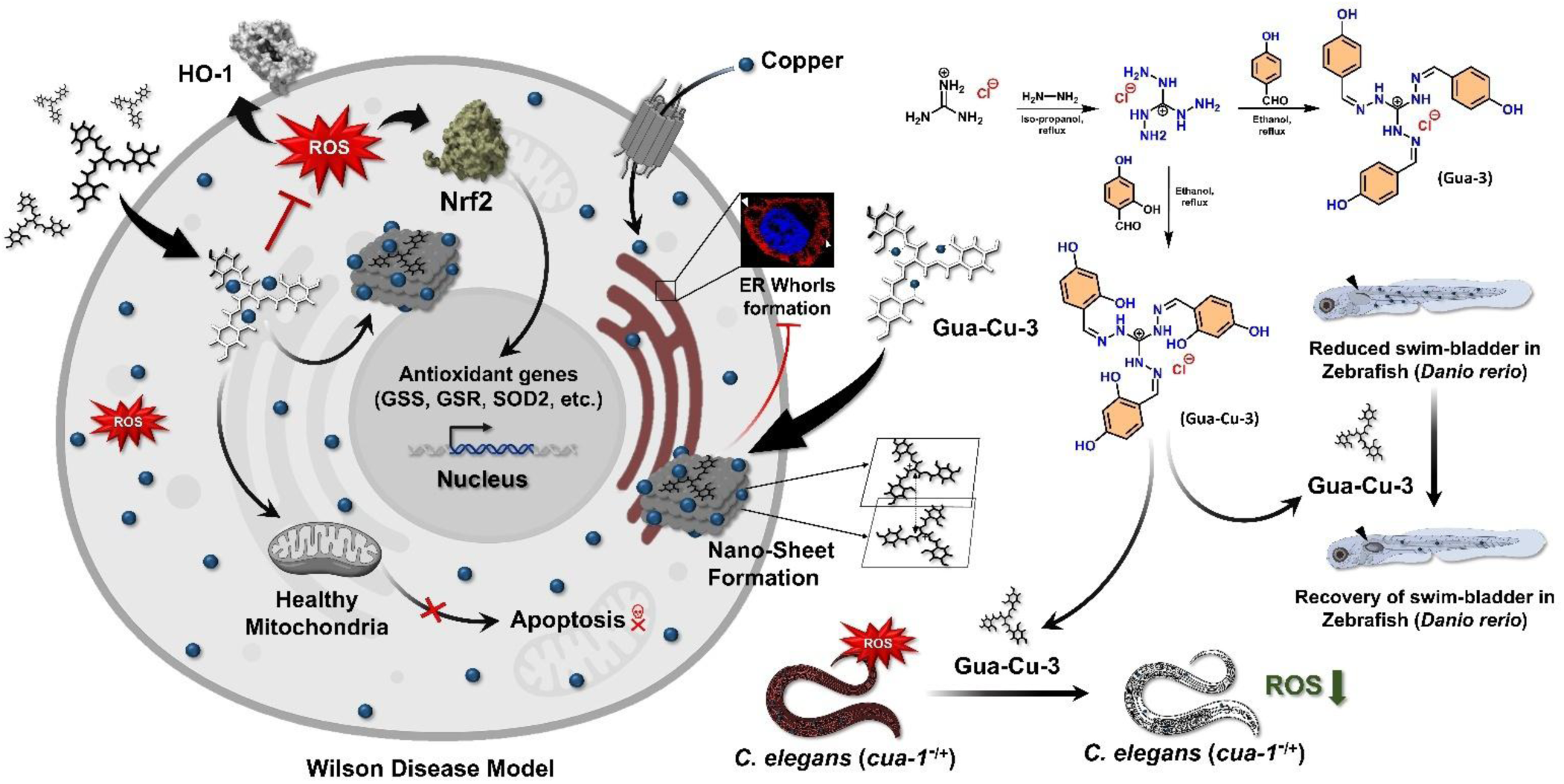
Proposed mechanism by which Gua-Cu-3 suppresses copper-driven oxidative stress and restores cellular homeostasis in Wilson disease models. Excess intracellular copper promotes redox cycling and reactive oxygen species (ROS) generation, leading to activation of oxidative stress pathways, including Nrf2/HO-1 signaling, endoplasmic reticulum stress i.e., ER Whorls formation, mitochondrial dysfunction, and apoptosis. Gua-Cu-3, a guanidinium-based copper-modulating scaffold, self-assembles into nanosheet-like architectures and coordinates labile copper pools, thereby suppressing copper-mediated ROS generation and downstream oxidative damage. This results in restoration of cellular homeostasis, improved mitochondrial integrity, reduced ER stress, and attenuation of apoptotic signaling. *In vivo*, Gua-Cu-3 reduces oxidative stress in *C. elegans* copper-homeostasis mutants and rescues copper-induced developmental defects in zebrafish, supporting its role as a dual-function modulator of pathological copper reactivity.

## Discussion

Copper dyshomeostasis presents a unique therapeutic challenge because pathology arises not only from the accumulation of excess copper but also from the oxidative chemistry mediated by redox-active labile copper species.^51^. Copper imbalance in Wilson’s disease causes oxidative stress, leading to increased lipid peroxidation, organelle dysfunction, and, ultimately, cell death. Currently used therapeutic options are mainly focused on reducing labile copper levels, either through chelation therapy or inhibition of intestinal copper absorption. While these treatments have been highly beneficial for patients, they have failed to address the oxidative damage caused by copper imbalance.^52^ Our finding is driven by the assumption that dual treatment targeting both labile copper levels and oxidative stress can be effective in dealing with copper-associated pathologies. Gua-Cu-3 has been successfully designed as a chemically defined scaffold with both copper binding and antioxidant properties. Multiple orthogonal techniques, including DFT calculations, UV-visible spectroscopy, isothermal titration calorimetry, NMR spectroscopy, EPR spectroscopy, and DOSY measurements, consistently support direct interaction between Gua-Cu-3 and copper ions. Importantly, the measured binding affinity is substantially lower than that typically associated with conventional high-affinity copper chelators, suggesting that Gua-Cu-3 engages bioavailable copper through a moderate-affinity interaction. While the precise intracellular copper species targeted by Gua-Cu-3 remain to be determined, this binding profile is consistent with a strategy aimed at modulating excess copper availability rather than indiscriminate removal of tightly bound physiological copper from cuproproteins which may not be the case for conventional high-affinity chelators that can strip copper from cuproproteins which leads to various complications. One of the key consequences of copper coordination by Gua-Cu-3 is the inhibition of copper-mediated oxidative chemistry more efficiently than non-chelating control Gua-3. In a variety of free radical assays, we found that Gua-Cu-3 inhibited the production of hydroxyl radicals and blocked copper-mediated oxidative chemistry, whereas the non-chelating control compound was less efficient. As can be observed from these results, the compound Gua-Cu-3 is not merely a radical scavenger but rather a chemical modulator of copper reactivity. The reason for this statement is very simple, inhibiting radical generation will work much better at preventing oxidative damage than inhibiting only the chain reaction involved in ROS production. Based on our findings, the influence of copper coordination in the cellular environment has been observed through a variety of experiments. In particular, the effect of Gua-Cu-3 was demonstrated in a cellular assay, where Gua-Cu-3 inhibited copper-dependent ATP7B trafficking from the TGN, helped recover Phen Green fluorescence, suppressed oxidative stress, decreased lipid peroxidation, and enhanced cellular viability. It should also be noted that Gua-3, which is structurally similar to Gua-Cu-3, showed antioxidative properties; however, in protecting against copper-induced oxidative damage, it was much less effective. This distinction suggests that antioxidant activity alone is insufficient to reverse wilson disease phenotype and highlights the importance of coupling redox protection with copper coordination. Moreover, the antioxidant activity of Gua-Cu-3 was evident from the normalization of numerous stress response pathways. Copper accumulation led to increased Nrf2 signaling, upregulation of HO-1 levels, loss of lysosomal integrity, mitochondrial superoxide generation, ER abnormalities, and apoptosis. All these were effectively reversed by the treatment with Gua-Cu-3. What must be highlighted here is that all experimental results can be explained by a single phenomenon: the sequestration of labile copper and the control of oxidative processes dependent on copper metabolism, thus averting a chain reaction of stress responses. Restoring lysosome, mitochondrial, and endoplasmic reticulum homeostasis, in particular, is important, as these organelle dysfunctions have recently emerged as key components of copper toxicity. Importantly, while the antioxidant effect of Gua-Cu-3 has been demonstrated using cell lines, it has also been shown in various biological systems, including *C. elegans* and Zebrafish. Such conservation of the activity of Gua-Cu-3 across species indicates that the chemical mechanism underpinning its copper sequestration and nd antioxidant action is organism-independent.The present study also revealed that Gua-Cu-3 undergoes supramolecular self-assembly into stable nanosheet-like architectures. Although the biological significance of this organization remains incompletely understood, the assemblies exhibited structural stability under physiologically relevant conditions. Whether supramolecular organization influences copper binding, cellular uptake, biodistribution, or biological activity warrants further investigation and represents an interesting avenue for future studies. Despite the encouraging results, several limitations must be considered. First, although Gua-Cu-3 exhibits selective copper binding and cytoprotection, the exact intracellular copper speciation and interaction with endogenous copper chaperones must still be determined^53^. Second, the long-term biological activity, pharmacokinetics, and systemic safety of Gua-Cu-3 have not yet been assessed in mammalian disease models^54^. Third, the contribution of supramolecular self-assembly to biological activity remains unresolved.^55,56^ Addressing these questions will be important for further development of the scaffold and for understanding how copper coordination and redox regulation can be optimally integrated within a single chemical framework. In general, this approach to chemical design could be applied to the development of metal-modulating drugs for other diseases in which other redox active metals, such as copper, iron, and redox stress, are involved, such as neurodegenerative disorders. In summary, this study identifies Gua-Cu-3 as a multifunctional copper-modulating scaffold capable of attenuating copper-associated oxidative stress and restoring cellular homeostasis across cellular and organismal models of copper dyshomeostasis. By integrating copper coordination with redox protection, Gua-Cu-3 provides a framework for the development of therapeutic strategies that address both metal imbalance and its downstream oxidative consequences.

## Materials and Methods

### ESI-MS

High-resolution Electrospray ionization mass spectrometry (HR-ESI-MS) was used to confirm the molecular masses of Gua-Cu-3 (MW 465 Da) and Gua-3(MW 417 Da) produced peaks consistent with their predicted molecular weight. Samples were dissolved in methanol and analyzed in positive-ion mode. The observed molecular ion peaks were consistent with the calculated molecular weights of Gua-Cu-3 (expected mass = 465 Da) and Gua-3 (expected mass = 417 Da), confirming successful synthesis of both compounds.

### ¹H NMR, ^13^C NMR and DOSY NMR titrations

NMR spectra were recorded using a Bruker spectrometer operating at the appropriate frequencies for ¹H and ¹³C nuclei. For ¹H NMR titration experiments, 8-10 mg Gua-Cu-3 was dissolved in 500 µL DMSO-d₆ and spectra were recorded. For copper-binding studies, CuCl₂ (1 Molar solution in DMSO-d₆; 35 µL) was mixed to 8-10 mg Gua-Cu-3 which was dissolved in 500 µL DMSO-d₆ and spectra were recorded after equilibration. Changes in chemical shifts were monitored to assess copper coordination.

### TEM /SEM

For self-assembly studies, Gua-Cu-3 and Gua-3 (1 mg each) were separately dispersed in 1 mL of 1X PBS buffer. The suspensions were sonicated for 45 min to ensure homogeneous dispersion and initiate self-assembly. After standing for 15 min at room temperature, aliquots of the solutions were drop-cast onto carbon-coated copper TEM grids for transmission electron microscopy (TEM), silicon wafers for field-emission scanning electron microscopy (FE-SEM). The samples were allowed to dry under ambient conditions before imaging studies. Morphological characterization was performed to evaluate nanosheet formation and supramolecular organization.

### Atomic Force Microscopy (AFM)

Atomic Force Microscopy (AFM) was employed to investigate the three-dimensional (3D) surface morphology of the synthesized nanosheets. AFM measurements were performed using an Oxford Instruments AFM system operating in tapping mode to minimize sample damage and achieve high-resolution topographical data. A small volume of the sample solution was drop-cast onto a clean silicon wafer substrate and allowed to dry under ambient conditions overnight to ensure uniform deposition. Surface analysis was conducted using AFM tips with a nominal radius of curvature appropriate for nano-scale imaging. Surface topography and morphology of the nanosheets were obtained by comparing the height and phase images taken at similar imaging conditions.

### Kelvin Probe Force Microscopy (KPFM) surface potential

Surface potential distribution and electronic properties were analyzed using Kelvin Probe Force Microscopy (KPFM). Solutions of the samples were cast as drops onto a silicon wafer substrate and left overnight at room temperature to achieve stable deposition. The AFM system from Oxford Instruments was used for measurements and capable of performing KPFM analysis with conductive Pt/Ir-coated silicon cantilevers to measure topography and potential simultaneously. The imaging in KPFM mode was performed using the tapping mode at an optimal lift height to avoid crosstalk between the morphological and electrical signals. The topography and surface potential images were collected in succession under identical scanning conditions. The surface potential was measured to determine the charge state of the self-assembled structures.

### Powder X-ray Diffraction (PXRD)

The PXRD experiments were carried out on a Rigaku benchtop diffractometer that utilizes a D/teX Ultra detector and Cu Kα radiation (λ = 1.5406 Å). Diffraction curves were recorded from 5 to 80 degrees 2θ at a scan speed of 2° min⁻¹ using the X-ray tube run at 40 kV. The obtained diffraction curves were then analyzed to assess crystallinity and lamellar ordering in the formed assemblies.

### HPLC analysis for Gua-Cu-3 stability

Gua-Cu-3 stability was assessed using reverse-phase HPLC (Waters system) using a Waters HPLC system equipped with a UV detector. Gua-Cu-3 (1 mg) was dissolved in 1 mL of methanol and analyzed using an isocratic acetonitrile: water (70:30, v/v) mobile phase on a C18 reverse-phase column. Chromatograms were acquired at start point(0 hour) and at 12 hour, monitoring UV absorbance at 278 nm. Peak retention time and chromatographic profile were compared to assess compound stability.

### Computational Methodology, Geometry Optimization and Binding Energetics

The ground state geometries of the ligand and the copper bound complex were optimized within the framework of Density Functional Theory (DFT) using the Gaussian 09 software package. Ground-state geometries of free Gua-Cu-3 and the corresponding Cu(II)-bound complex were subjected to full geometry optimization without any symmetry or structural constraints. The hybrid B3LYP (Becke three-parameter Lee-Yang-Parr) functional was employed for all calculations^57,58^. To balance computational efficiency and accuracy, the LanL2DZ effective core potential was utilized for the copper atoms, while the 6-31G+(d,p) split-valence double-zeta polarized and diffuse basis set was applied to all remaining atoms^59,60^. Solvent effect was considered in the calculation using an implicit water solvent model known as IEFPCM. As shown in the following equation, the interaction energy was calculated based on following formula.

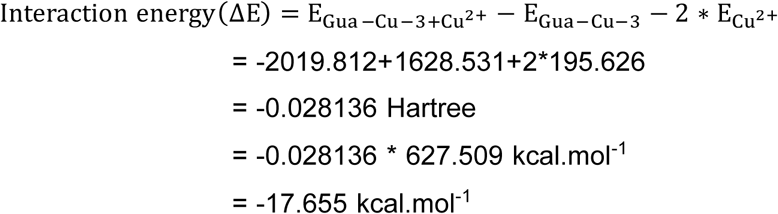

### Conceptual density functional theory (CD-DFT) Analysis

The conceptual density functional theory (CDFT) was used to analyze the electronic characteristics and reactivity of Gua-Cu-3 along with its Cu(II) complex. Important parameters such as the chemical potential (μ), chemical hardness (η), and electrophilicity index (ω) have been determined through HOMO and LUMO energies by using Koopman’s theorem.^61,62^

As the copper atom is bind to the ligand molecule, there is a significant reduction in the HOMO-LUMO energy gap. It suggests that the polarizability and electron delocalization of the complex have been enhanced. These changes are consistent with metal-ligand charge transfer interactions and the experimentally observed bathochromic shift in UV-vis spectra.

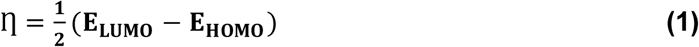

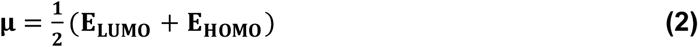

The global electrophilicity index (**ꞷ**) is given as

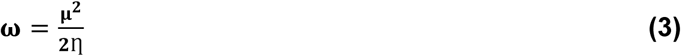

Where, **E_LUMO_** and **E_HOMO_** are energies of the lowest unoccupied molecular orbital (LUMO) and highest occupied molecular orbital (HOMO) respectively^63^.

### UV-vis spectroscopy

UV-visible absorption measurements were performed using a Cary UV-visible spectrophotometer at room temperature. Gua-Cu-3 was dissolved in Milli-Q water and absorption spectra were recorded over the wavelength range of 200-600 nm. For metal-binding studies, CuCl₂ was added incrementally to the ligand solution and spectra were recorded after each addition following equilibration. All measurements were performed under identical instrumental settings and solvent conditions. Binding constant was calculated by Bind-fit software.

### Isothermal titration calorimetry (ITC)

ITC experiments were performed on a Malvern MicroCal analytical calorimeter at room temperature using 19 sequential injections spaced at 180 s intervals. The syringe was loaded with 1.0 mM CuCl₂, and the sample cell contained 12.5 µM Gua-3 or Gua-Cu-3. Raw thermograms and integrated heats were used to evaluate binding-associated heat changes.

### Electron paramagnetic resonance (EPR) spectroscopy

Production of hydroxyl radicals (•OH) was assessed through EPR spectroscopy employing 5,5-dimethyl-1-pyrroline N-oxide (DMPO) as the spin trap. Mixtures of Cu²⁺ and hydrogen peroxide with or without Gua-Cu-3 were added into quartz capillaries for EPR analysis.

EPR spectra were performed with a Bruker EMXmicro spectrometer operated at X-band frequency (9.32 GHz) at room temperature, where the magnetic field was modulated at 100 kHz while the microwave power was set to 48.83 mW and modulation amplitude of 10 G.

### Cell lines and cell culture

Human hepatocellular carcinoma cells (HepG2) were cultured in low-glucose Minimum Essential Medium (MEM) (Himedia, AL047S), supplemented with 10% fetal bovine serum (FBS) (Gibco, 10270-106) and 1× Pen/Strep (Gibco, 15140-122). The cells were grown at 37°C in an incubator with a humidified environment having 5% CO₂, with cells being passaged periodically for their logarithmic growth. For all experiments, cells were used at ∼60-70% confluency unless specified otherwise.

### MTT assay for cell mortality assessment

Cell mortality was measured using MTT assay, which reflects NAD(P)H-dependent oxidoreductase activity that reduces MTT to insoluble purple formazan crystals. HepG2 cells were seeded into flat-bottom 96-well plates at 5000 cells per well in 200 µL complete MEM and incubated for 72 h at 37 °C, 5% CO₂ to allow attachment and growth till cell become 50-60% confluent. Cells were treated according to the experimental design and incubated for 24 h.

Following treatment, culture medium was gently removed, and MTT solution (prepared in PBS) was mixed with MEM at a final concentration of 0.1 mg/mL. Plates were incubated for 4 h at 37 °C in 5% CO₂. The medium was then removed, and formazan crystals were dissolved in 200 µL dimethyl sulfoxide (DMSO). Absorbance was measured at 570 nm using a multi-well plate reader. Cell viability was calculated relative to untreated controls.

### Fixed-cell immunofluorescence staining

After treatment completion, the media was discarded and washed twice with cold PBS. Cells were then fixed in 2% paraformaldehyde (PFA) in PBS for 20 min, followed by quenching with 50 mM NH₄Cl for 20 min. Cells were blocked with 3% BSA in permeabilization buffer (PBSS: 0.075% saponin in PBS) for 1.5 h. Primary antibodies were incubated for 1.5 h at room temperature, followed by secondary antibodies for 40 min. Cells were washed with PBSS (3×) and PBS (2×), then mounted with Fluoroshield with DAPI (Sigma, #F6057). Antibodies used: ATP7B (Abcam #ab124973, 1:400), Golgin-97 (CST #13192, 1:400), Nrf2 (Affinity #AF0639, 1:200), Calnexin (Abcam #ab22595.)

### Phen Green FL live cell imaging for determination of labile copper pool

To determine the labile copper pool, live-cell imaging was performed. Cells were seeded in confocal imaging dishes (SPL Lifesciences, #200350). Following drug treatment, cells were stained with 10 µM Phen Green FL in serum-free MEM for 30 min. During microscopy, cells were maintained in phenol red-free, low-glucose MEM imaging medium (Gibco, #41500-034) containing 2% FBS, 20 mM HEPES, and 1% Trolox as antifading agent(Sigma, #238813). Fluorescence images were acquired using a Leica SP8 confocal microscope(63× oil immersion objective,NA 1.4).

### DPPH radical scavenging assay

Antioxidant activity was evaluated using the DPPH radical scavenging assay. Briefly, freshly prepared methanolic solution of DPPH (200 µM) was mixed with varying concentrations of Gua-Cu-3 or Gua-3 in a 96-well plate. Following 30 minutes incubation in the dark at room temperature for the designated period, absorbance was measured at 517 nm using a BioTek Cytation 5 multimode plate reader. Ascorbic acid was included as a positive control. Radical scavenging activity was expressed as the percentage reduction in DPPH absorbance relative to untreated controls.

### ABTS assay

The antioxidant activity of Gua-Cu-3 and Gua-3 was also evaluated using the ABTS [2,2′-azinobis (3-ethylbenzothiazoline-6-sulfonic acid)] radical cation decolorization assay. The ABTS radical cation (ABTS•⁺) was generated by reacting ABTS (7 mM) with potassium persulfate (2.45 mM) in distilled water and allowing the mixture to stand in the dark at room temperature for 12-16 h prior to use. Before the assay, the ABTS•⁺ solution was diluted with phosphate-buffered saline (PBS) or ethanol to obtain an absorbance of approximately 0.70 ± 0.02 at 734 nm. In a 96-well plate, varying concentrations of Gua-Cu-3 or Gua-3 were mixed with the ABTS•⁺ working solution. After incubation for 10 min at room temperature in the dark, absorbance was recorded at 734 nm using a BioTek Cytation 5 multimode plate reader. Ascorbic acid was used as a positive control. The percentage inhibition of ABTS radical cation can be calculated by using:

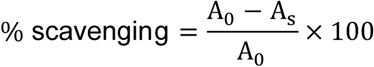

where A_0_represents the absorbance of the ABTS•⁺ solution without antioxidant and A_s_represents the absorbance in the presence of the test compound.

### Deoxyribose Degradation Assay (Hydroxyl Radical Scavenging)

Hydroxyl radical-mediated oxidative damage was evaluated using the deoxyribose degradation assay. Reaction mixtures (1 mL total volume) contained 2-deoxy-D-ribose (2.8 mM), phosphate buffer (20 mM, pH 7.4), CuCl₂, H₂O₂ (1 mM), and ascorbic acid (0.1 mM). Various concentrations of Gua-Cu-3 or Gua-3 were added and the reactions were incubated at 37 °C for 1 h. Following incubation, 1 mL of thiobarbituric acid (TBA, 1% w/v) and 1 mL of trichloroacetic acid (TCA, 2.8% w/v) were added. Samples were heated at 95 °C for 15 min, cooled to room temperature, and absorbance was measured at 532 nm using a BioTek Cytation 5 multimode plate reader. Oxidative degradation of deoxyribose was quantified relative to untreated controls.

### Zebrafish maintenance, breeding, and embryo handling

Experiments involving zebrafish were carried out using the *Danio rerio* Tübingen strain. The adult fish were grown in a recirculating water system maintained at 28 °C, and light/dark cycles were maintained at 14 h and 10 h respectively. The fishes were provided with dried foods twice daily (9 A.M and 5 P.m.) and alive brine shrimp (Artemia sp.) once daily (2 PM). In the case of embryo harvesting, the fish were housed in dedicated breeding tanks. Males and females’ fish were introduced together into the breeding tank and isolated by dividing the tank physically overnight. The barrier between them was removed the following morning, which allowed them to mate. After 1-hour, adult fish were isolated from the tank, eggs were harvested using a fine strainer, washed and transferred to embryo medium (5 mM NaCl, 0.17 mM KCl, 0.33 mM CaCl₂, 0.33 mM MgSO₄, 0.0001% methylene blue). The embryos were maintained at 28.5 °C. Exposure of the embryos to copper was done using the indicated CuCl₂ solution for 45 min at 28.5 °C.

### *C. elegans* strains and culture conditions

Wild-type *Caenorhabditis elegans* Bristol N2 and the copper-ATPase mutant strain cua-1 (tm12763) were used in this study. The cua-1(tm12763) strain was obtained from the National Bioresource Project (Mitani lab, Tokyo, Japan). Worms were maintained on nematode growth media (NGM) agar plates seeded with Escherichia coli OP50 as a food source. Worms were cultured at 20 °C, and synchronized populations were prepared as required for experiments. Animals were grown till L4 larval stage, which was further used for each experiments. For compound treatment, synchronized L4 worms were treated with Gua-Cu-3 overnight under the indicated experimental conditions. Following treatment, worms were then washed, anesthetized using 0.5 mM sodium azide, fixed on 0.3% agar pads, and imaged.

### Measurement of malondialdehyde (MDA) using the TBARS assay

Lipid peroxidation was quantified by measuring malondialdehyde (MDA) using a commercial colorimetric TBARS assay kit (BioVision, Milpitas, CA, USA) according to the manufacturer’s instructions. Following the indicated treatments, Cells were washed with PBS, trypsinized, and centrifuged. Pellets were processed according to the kit instructions. In brief, lysates were prepared and centrifuged, and 200 µL of supernatant was mixed with 600 µL thiobarbituric acid (TBA) reagent and incubated at 95 °C for 1 h to form the MDA-TBA adduct. Samples were cooled to room temperature, and 200 μL aliquots were transferred to a 96-well plate, and absorbance was measured at 532 nm using a UV-Vis spectrophotometer.

### Immunoblotting

Following CuCl₂ and Gua-Cu-3 treatments, cells were pelleted and lysed in 120 µL RIPA buffer (10 mM Tris-Cl, pH 8.0, 1.0% Triton X-100, 1 mM EDTA, 0.5 mM EGTA, 0.1% sodium deoxycholate, 0.1% SDS, 140 mM NaCl) supplemented with 1× protease inhibitor cocktail and 1 mM PMSF. Lysates were incubated on ice for 30 min and probe-sonicated (6 pulses; 5 s ON, 25 s OFF; 100 mA). Protein concentration was determined using Bradford reagent (Merck, #B6916). Equal amounts of protein (20 µg) were mixed with 4× NuPAGE loading buffer (Invitrogen, #NP0007) to a final 1× concentration and resolved on 12% SDS-PAGE gels. Proteins were transferred to nitrocellulose membranes (Millipore, #IPVH00010) using semi-dry transfer. Membranes were blocked in 3% skim milk in TBST (TBS with 0.1% Tween-20) for 3 h at room temperature with gentle shaking. Primary antibodies were incubated overnight at 4 °C, followed by washes in TBST (0.01% Tween-20). Secondary antibody incubation (anti-rabbit HRP; CST #7074; 1:6000) was performed for 1.5 h at room temperature. Membranes were washed (TBST 3×, TBS 2×) and developed using Clarity Max ECL substrate (Bio-Rad, #1705062). Chemiluminescence was recorded on a ChemiDoc system (Bio-Rad). Primary antibodies and dilutions: α-tubulin (Affinity #AF7010, 1:20000), HO-1 (Affinity #AF5393, 1:1500), LAMP1 (1:400; DSHB). Band intensities were quantified using ImageJ software and normalized to loading control, α-tubulin.

### CellROX-based oxidative stress assay in HepG2 cells (flow cytometry)

Oxidative stress was evaluated utilizing the fluorescent dye CellROX Green. CellROX Green is a fluorometric probe, which upon oxidation by free radicals becomes highly fluorescent (λex/λem = 508/525 nm). The HepG2 cells were grown in 24-well plates and pretreated with Gua-Cu-3 for 3 hours, after which the cells were exposed to CuCl₂. After 24-hour incubation, the treated cells were stained with CellROX Green at 5 µM concentration for 20-30 minutes.

Cells were washed with DPBS 1× and dissociated by trypsinization using 100 µL Trypsin-EDTA 0.25% (Gibco, #25200072). Dissociation was followed by addition of 300 µL of FACS buffer (DPBS 1×, 2% FBS, 25 mM HEPES, 2 mM EDTA) and then single cell suspension was prepared. Analysis of samples was carried out using BD LSR Fortessa flow cytometer (BD Biosciences). 10,000 events were collected for each sample, and the mean fluorescence intensity (MFI) was used for quantification.

### CellROX live-cell imaging in HepG2 cells (confocal microscopy)

The confocal microscopy of the intracellular level of ROS was performed with cells growing on confocal dishes (SPL Lifesciences, #200350). The HepG2 cells were stained with CellROX Green (5 μM) and then washed before the image acquisition procedure. For imaging, the cells were maintained in the imaging medium consisting of low glucose MEM media without phenol red (Gibco, #41500-034) with 2% FBS, 20 mM HEPES, and 1% Trolox used as an antifading agent (Sigma, #238813). Confocal imaging was conducted using a Leica SP8 microscope equipped with a 63× oil immersion objective lens with NA = 1.4. Images were processed using Leica Lightning deconvolution software, and fluorescence intensity was quantified using ImageJ.

### Oxidative stress in zebrafish embryos (CellROX Orange)

To measure ROS in zebrafish embryos, embryos were kept in E3 medium and pre-treated with Gua-Cu-3(25 µM) for 30 min in E3 media. Embryos were then washed with E3 and exposed to copper (200 µM) in fresh E3 medium. After treatment, embryos were incubated with 20 µM CellROX Orange for 30 min. Embryos were anesthetized with 0.003% tricaine (Sigma, #A-5040) in E3 medium and single-cell suspensions were prepared by incubating the embryo in dissociation solution (Collagenase 1mg/ml in HBSS with Phenol Red, Trypsin-EDTA, Tricaine, HBSS without Ca^2+^,Mg^2+^) for 30 Minutes and subsequent Pipette up and down 20 times in interval of 5 minutes to facilitate tissue dissociation for 4 times and Pass homogeneous cell suspension through 70 μm strainer and mean/median fluorescence intensity was recorded using BS LSR Fortessa.

### Oxidative stress in *C. elegans* (CellROX Orange)

ROS levels in worms were quantified under basal conditions and after a 12 h treatment with Gua-Cu-3 (25 µM). CellROX Orange (Invitrogen) was used diluted in DMSO and used at a final concentration of 10 µM for staining worms. Worms were washed thrice with M9 buffer to eliminate OP50 bacteria, which act as the food material for *C. elegans*; 100 µL of the M9 buffer containing worm was mixed with 100 µL CellROX solution (10 µM in M9) and incubated for 1 h at 20 °C in the dark. Worms were fixed on 3% agar pads containing sodium azide and imaging was done using a Leica TCS SP8 confocal microscope.

### Determination of mitochondrial superoxide using MitoSOX Red

Superoxide radical from mitochondria was measured using Mitosox Red (Invitrogen; Cat. No.: M36008). Mitosox Red is a fluorescence stain capable of identifying superoxide radicals within mitochondria. Within the mitochondria, the Mitosox reagent gets oxidized by the superoxide radicals, which results in the production of fluorescence signals that can be quantitated for superoxide radicals within the mitochondria. Following Gua-Cu-3 and Cu treatment, the cells were treated with 2.5 µM Mitosox Red for 30 minutes, followed by washing with DPBS and trypsinization of the cells. The cells were suspended in 350 µL of FACS buffer, and data acquisition was performed using BD LSRFortessa flow cytometer.

### Annexin V/PI staining and flow cytometry

Apoptosis in HepG2 cells was evaluated using Annexin V Alexa Fluor 488 and propidium iodide (PI). Cells were grown in 12-well plates and pretreatment was done with 25 µM Gua-Cu-3 for 3 h, then treated with 500 µM CuCl₂ for 24 h. After treatment completion, cells were washed two times with PBS, detached with 0.25% Trypsin-EDTA, and pelleted (38 × g, 2 min). Pellets were washed twice in PBS (without Ca²⁺/Mg²⁺) and resuspended in diluted (4 times in MiliQ water) Annexin V binding buffer. 5 µL Annexin V Alexa Fluor 488 (Invitrogen, A13201) was added in cell suspension according to the manufacturer’s instructions, and samples were incubated for 30-40 min at room temperature in the dark.

Propidium Iodide (Sigma, #P-4864-10ML) was diluted at 1:10 ratio in Annexin V binding buffer, and 4 µl were added to each sample at a final concentration of 2 µg/ml. Incubation was allowed to proceed for 5 minutes at RT, followed by washing with 500 ul binding buffer, centrifuging, and resuspension in 350 µl FACS buffer for analysis of apoptotic cells. Simultaneously, single cell suspension from 3 dpf zebrafish embryos was prepared and analyzed by Annexin V/PI following the same procedure.

### Image analysis and statistics

Image analysis was performed using ImageJ/Fiji. For colocalization studies, Pearson’s correlation coefficient (PCC) was calculated using the Colocalization Finder plugin. Regions of interest (ROIs) were manually selected from optimal z-stacks for each cell. Pearson correlation coefficient (PCC) was used to quantify colocalization. For ATP7B trans-Golgi network (TGN) colocalization with Golgin-97, the fraction of ATP7B signal overlapping Golgin-97-marked compartments was quantified using PCC. Nuclear localization of Nrf2 was quantified by splitting RGB channels and manually outlining nuclear ROIs. Where not otherwise mentioned, all experiments were conducted with at least three independent biological replicates. Box-and-whisker plot shows median and 25%-75% percentile. Whiskers are drawn to values which are ±1.5 times of IQR. In case of comparison between two unpaired groups, non-parametric test was applied (Mann-Whitney U test). Statistical significance was considered as *p<0.05, **p<0.01, ***p<0.001, and ****p<0.0001. GraphPad prism software was utilized for statistical calculations and graph construction. The macro script for ImageJ can be found on GitHub.

### Animal ethics statement

The zebrafish were cultured in the recirculating aquaculture system, and breeding was regularly carried out using breeding tanks. Experiments were carried out following the guidelines for conducting research as per the Institutional Animal Ethics Committee of the Indian Institute of Science Education and Research, Kolkata. All procedures performed involving *C. elegans* were performed in accordance with the Guidelines on Handling and Training of Laboratory Animals formulated by the Indian Institute of Science Education and Research, Kolkata.

### Safety

There are no unexpected, new, and/or significant hazards or risks associated with this work.

## Acknowledgement

This work was supported by DBT-Wellcome Trust India Alliance Fellowship (IA/I/16/1/502369), STARS-2 Grant (2023-0210) from Ministry of Education, Govt. of India, Department of Science and Technology (Therapeutic strategies for prevalent Rare/Orphan Disorders; DST/TDT/TC/RARE/2022/32), Govt. of India and IISER K intramural funding to A.G. A.D. acknowledges the ANRF-J.C. Bose Fellowship and funding through the JBR/2023/000005 grant. Zebrafish studies were supported by ANRF, New Delhi (CRG/2023/001075) and by DBT-Wellcome Trust India Alliance Fellowship (IA/I/18/2/504016) to C.P.. R.P. is supported by a pre-doctoral fellowship from the Department of Biotechnology, Government of India, India. A.N.R. is supported through Indian Council of Medical research grant (2012-9973) to A.G. S.S, D.B., A.J is supported by a pre-doctoral fellowship from the University Grant Commission, Govt of India. K.D. is supported by a predoctoral fellowship from IISER Kolkata. The authors also acknowledge the IISER Kolkata Central Instrumentation Facility (iCIF) for support and instrumental facilities.

## Author contributions (CrediT)

RP: Investigation, Methodology, Writing - original draft

ANR: Investigation, Methodology, Writing - original draft

SS: Investigation, Methodology

KD: Investigation, Methodology

DB: Investigation, Methodology

TG: Resources

AJ: Software, Investigation

KG: Conceptualization

CP: Resources

AD: Conceptualization, Project administration, Supervision, Writing - original draft

AG: Conceptualization, Project administration, Supervision, Writing - original draft

## Data availability statement

The data supporting this article have been included as part of the Supplementary Information.

## Supplementary data

**S1.**
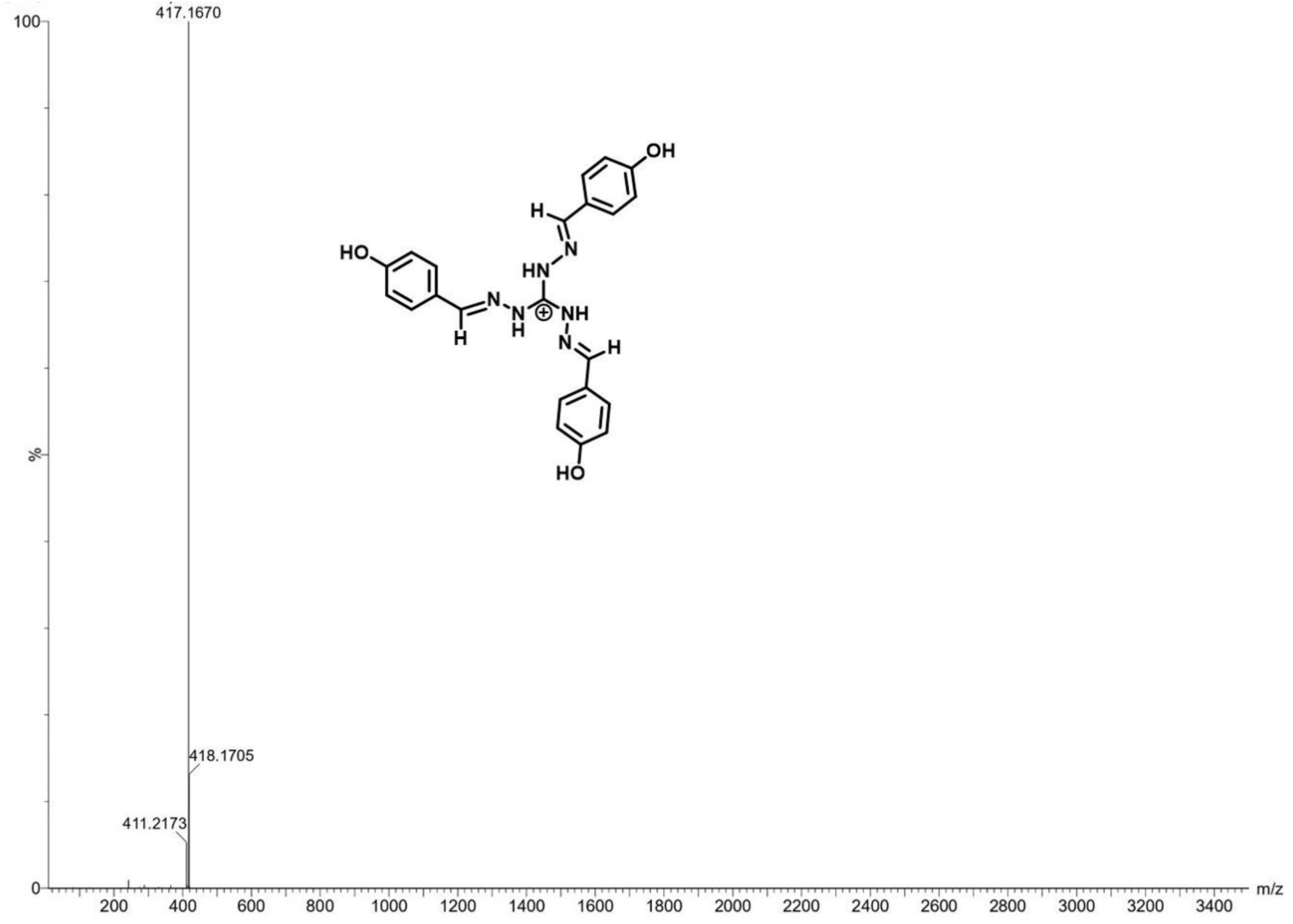
Ectrospray ionization mass spectrometry (ESI-MS) spectrum of the Gua-3. The spectrum recorded in positive ion mode shows a prominent molecular ion peak at m/z 417.1670, corresponding to the protonated species [M+H] ⁺ ion, consistent with the expected molecular formula of Gua-3. Minor peaks at m/z 418.1705 and 411.2173 correspond to natural isotopic distribution and fragment ions, respectively. The inset shows the proposed chemical structure of Gua-3.

**S2.**
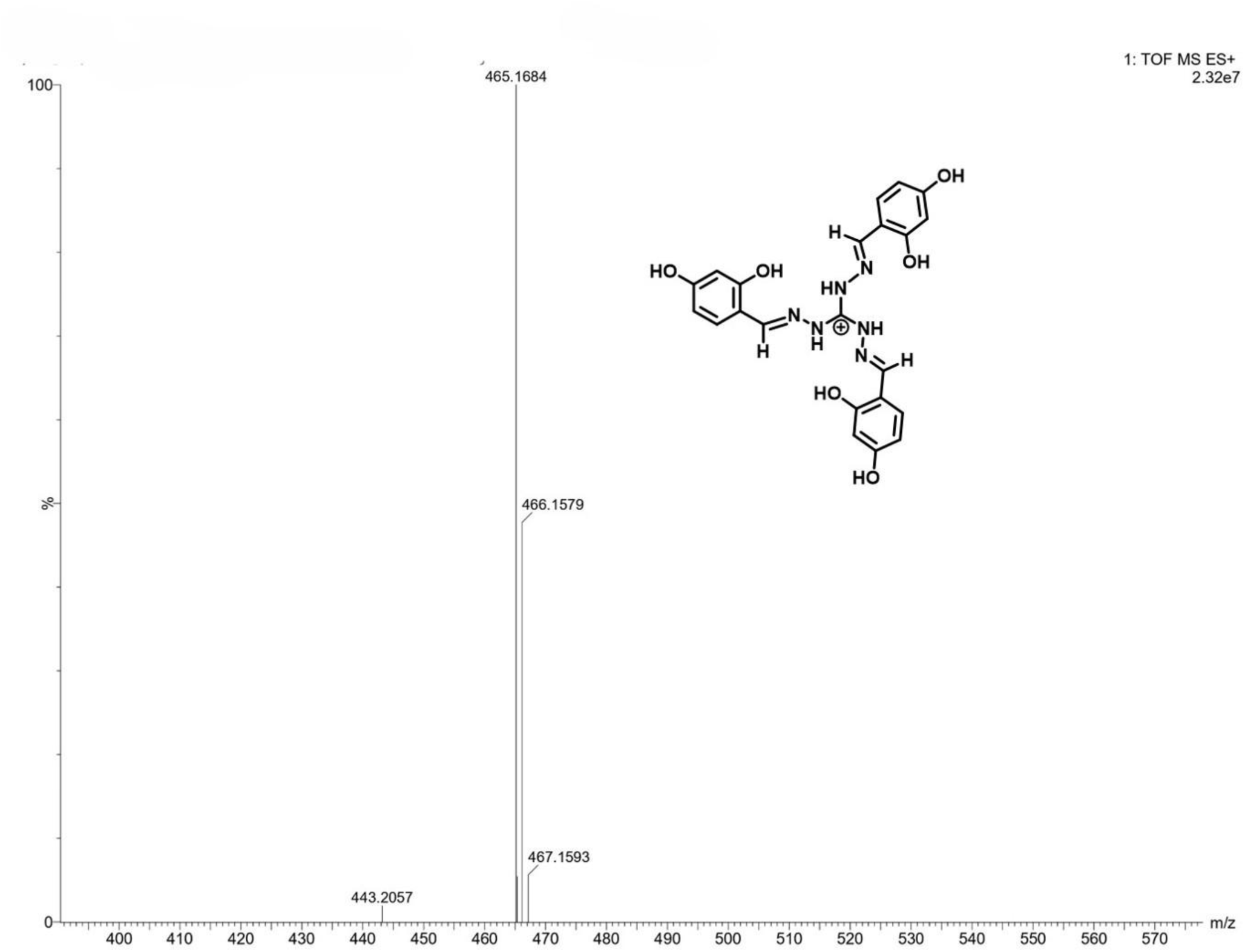
Electrospray ionization mass spectrometry (ESI-MS) spectrum of the Gua-Cu-3. The spectrum recorded in positive ion mode exhibits a prominent molecular ion peak at m/z 465.1684, corresponding to the protonated molecular ion [M+H] ⁺ ion, consistent with the expected molecular formula of Gua-Cu-3. The adjacent peaks at m/z 466.1579 and 467.1593 correspond to the natural isotopic distribution, while the A low-intensity signal at m/z 443.2057 is likely attributable to minor in-source fragmentation. The inset shows the proposed chemical structure of Gua-Cu-3.

**S3.**
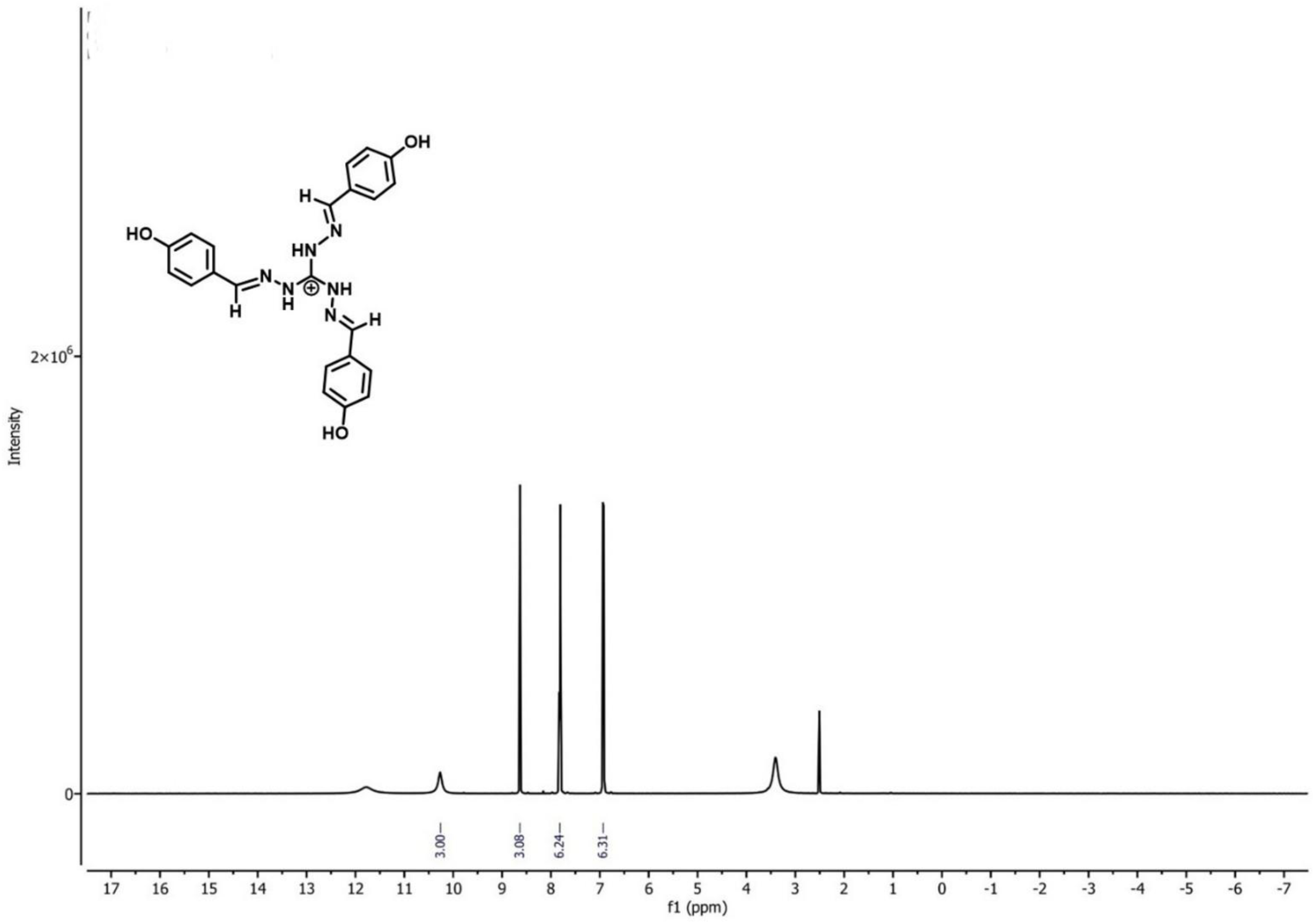
The spectrum displays resonances consistent with the proposed structure of Gua-3, including characteristic azomethine (-HC=N-) proton signals in the downfield region (δ ≈ 8.72 ppm) and aromatic proton resonances between δ ≈ 6.8-8.00 ppm. The integrations at peaks match the predicted proton stoichiometry for the molecule. The peak appearing at δ ≈ 3.3 ppm is attributed to residual water present in DMSO-d₆. Based on the information from the spectra, it appears that there was formation of the proposed guanidinium scaffold depicted in the inset figure.

**S4.**
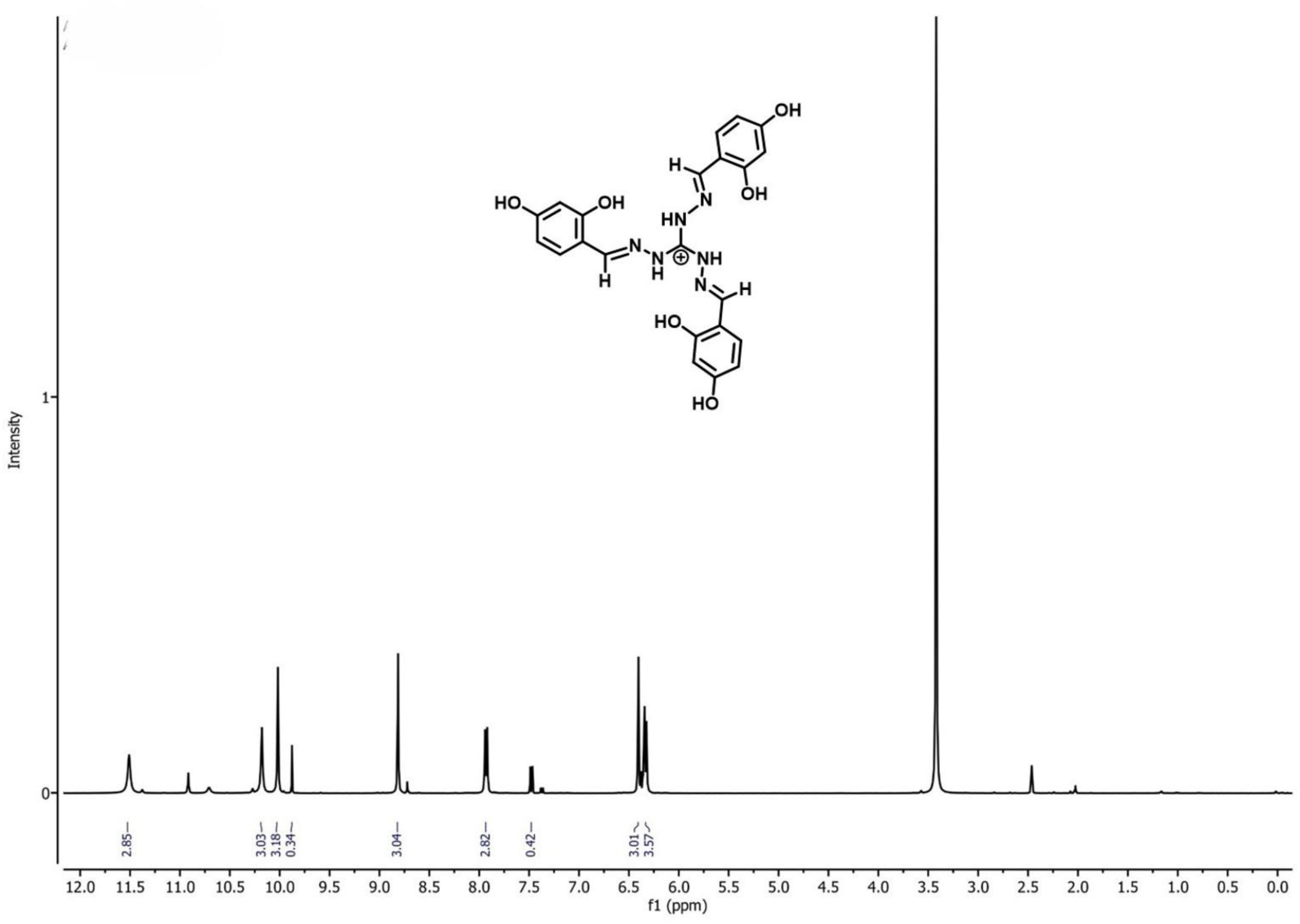
¹H NMR spectrum of Gua-Cu-3 (400 MHz, DMSO-d₆). These include the presence of downfield resonances of hydroxyl protons of phenol (δ≈ 8.7 ppm), HC=N protons (δ≈ 8.7 ppm) along with aromatic protons in the range δ≈ 6.0-8.5 ppm as suggested by the proposed structure of Gua-Cu-3. Peak integrations agree with the expected proton stoichiometry of the molecule. Signals at δ ≈ 3.33 ppm and δ ≈ 2.50 ppm correspond to residual water and DMSO-d₆, respectively. Collectively, the spectral features are consistent with formation of the proposed guanidinium-based scaffold shown in the inset.

**S5.**
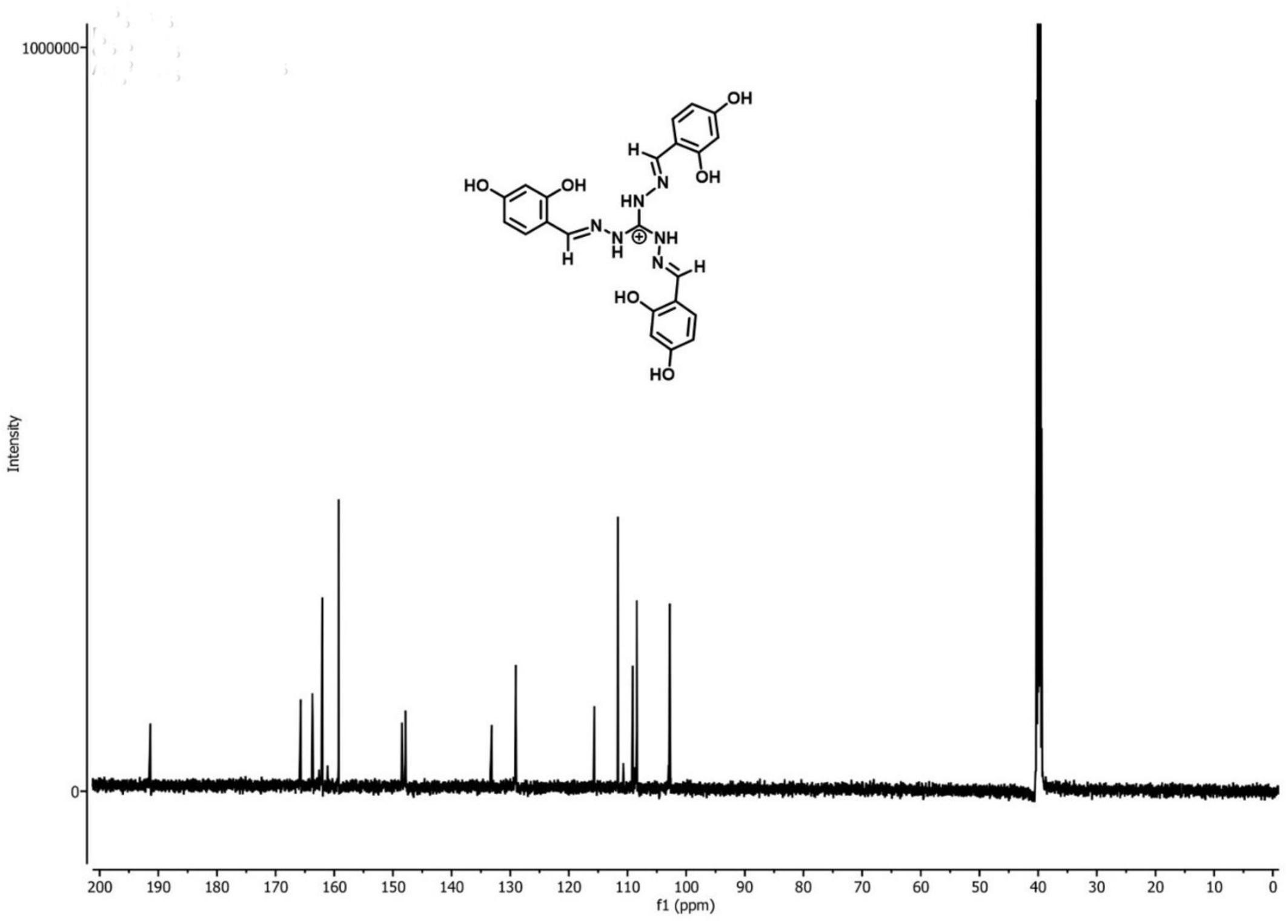
¹³C NMR spectrum of Gua-Cu-3 (100 MHz, DMSO-d₆). The spectrum exhibits resonances consistent with the proposed structure of Gua-Cu-3, including downfield signals in the δ ≈ 150-165 ppm region attributable to imine- and phenolic oxygen-substituted carbons, together with aromatic carbon resonances between δ ≈ 100 and 150 ppm. The reduced number of observed resonances relative to the total carbon count is consistent with the molecular symmetry of the molecule. The signal at δ ≈ 39.5 ppm corresponds to residual DMSO-d₆. Overall, the observed carbon resonances are consistent with formation of the proposed guanidinium-based scaffold shown in the inset.

**Figure S6.**
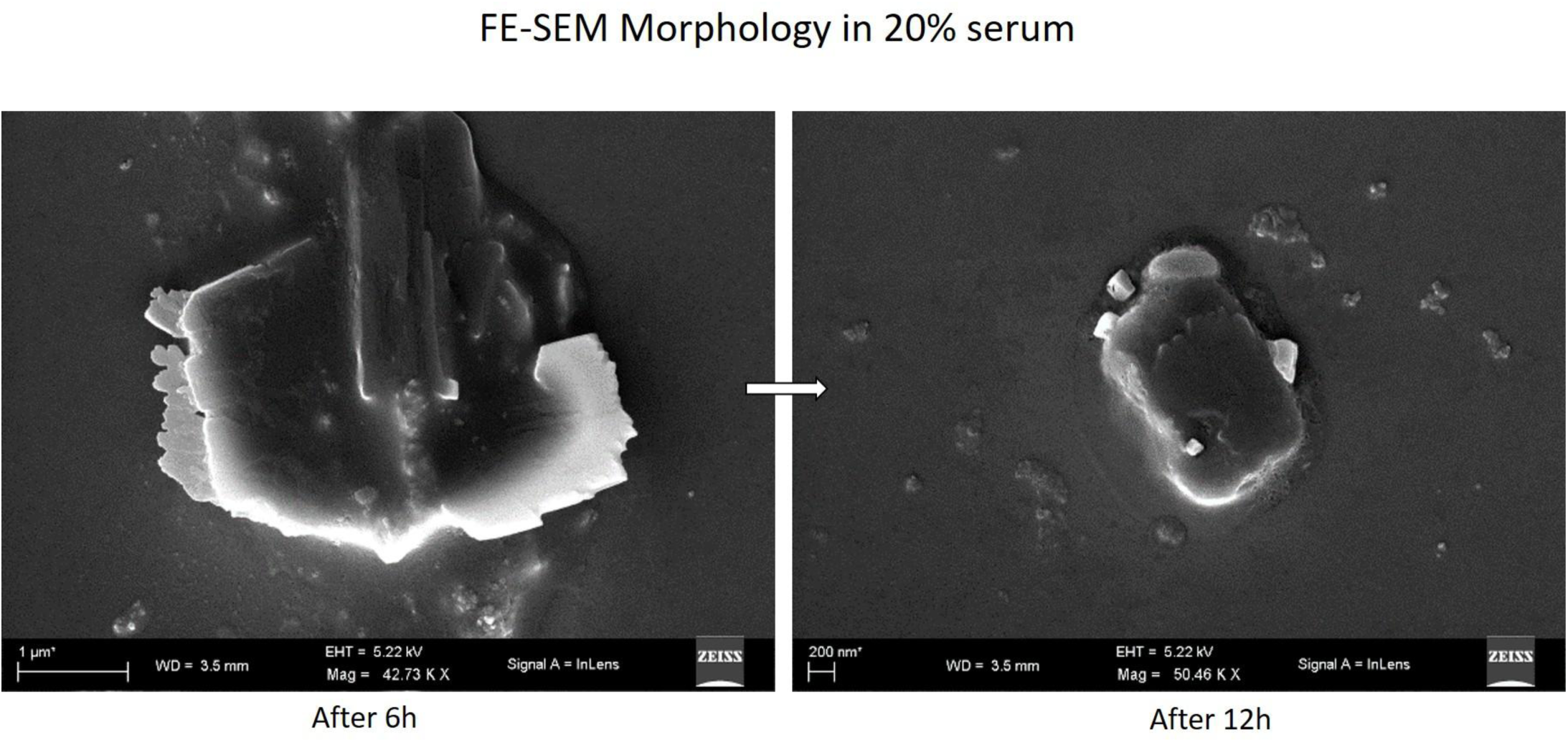
Representative FE-SEM images of Gua-Cu-3 following incubation in 20% serum-containing medium. Sheet-like supramolecular assemblies were observed after 6 h and remained detectable after 12 h of incubation, indicating preservation of the overall nanosheet morphology under serum-containing conditions. Images are representative of multiple examined fields. Scale bars are as indicated.

**S7.**
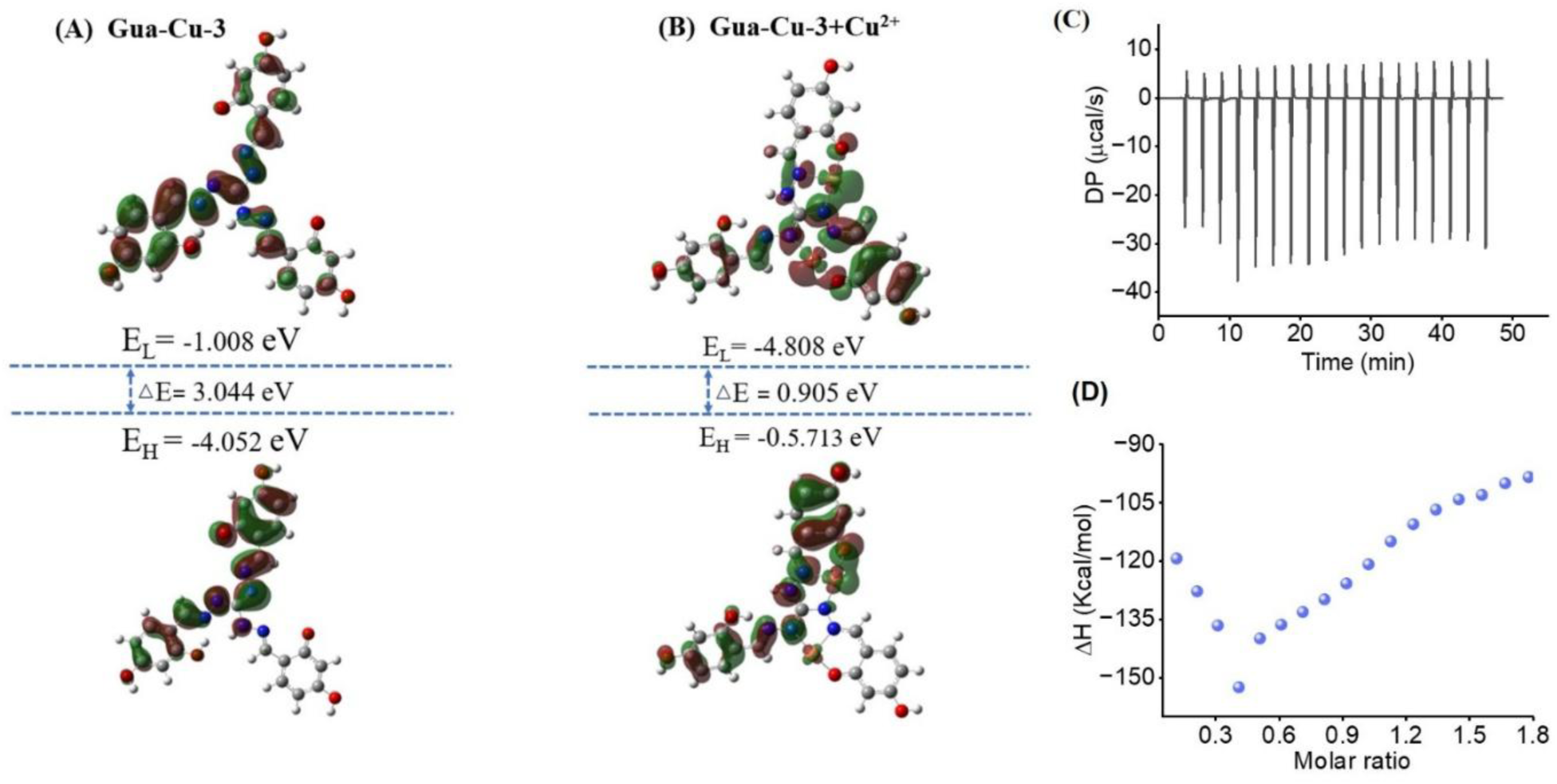
(A, B) Frontier molecular orbital (HOMO and LUMO) distributions and energy levels of free Gua-Cu-3 (A) and the Cu(II)-bound Gua-Cu-3 complex (B) calculated at the B3LYP/6-31G+(d,p)/LanL2DZ level. Copper coordination markedly reduces the HOMO-LUMO energy gap (ΔE), indicating enhanced electronic delocalization and altered electronic properties of the complex. (C) Representative ITC thermogram obtained from titration of Cu(II) into Gua-3 under identical experimental conditions used for Gua-Cu-3. Minimal heat changes were observed throughout the titration. (D) Integrated heat profile corresponding to panel C. The absence of a saturable binding isotherm indicates no measurable interaction between Gua-3 and Cu(II) under the experimental conditions employed.

**S8.**
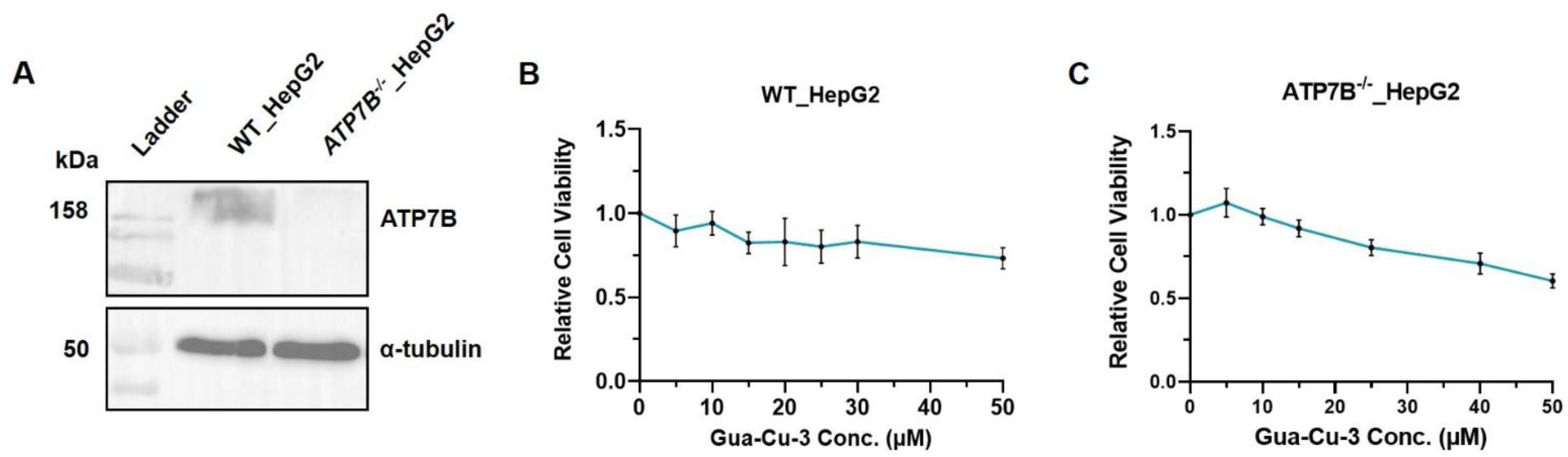
Validation of ATP7B knockout and cytotoxicity assessment of Gua-Cu-3 in HepG2 cellular models. (A) Immunoblot analysis confirming successful deletion of ATP7B in CRISPR-Cas9-generated ATP7B^⁻/⁻^_HepG2 cells. ATP7B protein is detected in WT HepG2 cells but is absent in ATP7B^⁻/⁻^_HepG2 cells. α-Tubulin serves as a loading control. (B, C) Cell viability analysis of WT HepG2 (B) and ATP7B^⁻/⁻^_HepG2 (C) cells after exposure to increased concentrations of Gua-Cu-3 for 24 h using MTT assay. There was little toxicity induced by Gua-Cu-3. Highest concentration used for Gua-Cu-3 is 25 μM in this study, indicating that it is appropriate for use and non-toxic.

**S9.**
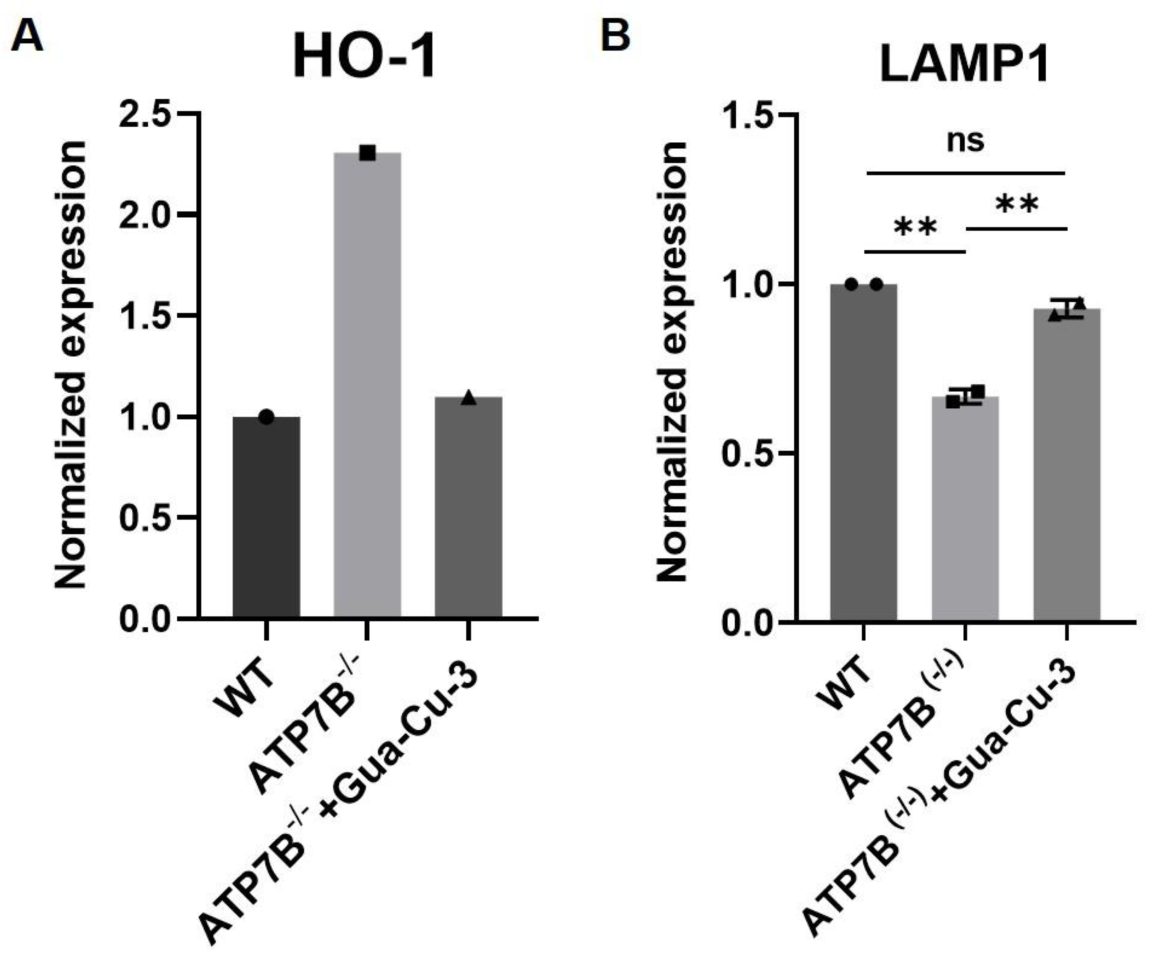
(A) Densitometric analysis of HO-1 protein levels normalized to α-tubulin. ATP7B deficiency increased HO-1 expression compared to wild-type cells, in line with increased oxidative stress. Gua-Cu-3 treatment decreased HO-1 expression towards basal level. **(B)** Densitometric analysis of LAMP1 protein levels normalized to α-tubulin. ATP7B deficiency led to a decrease in LAMP1 expression, suggesting impaired lysosomes, while Gua-Cu-3 treatment increased LAMP1 expression to the level of wild-type cells. All experiments were normalized to wild-type control cells using α-tubulin as the loading control.

